# A reinforcement learning framework for goal-directed stimulation of engineered neuronal networks

**DOI:** 10.64898/2026.04.14.718383

**Authors:** Benedikt Maurer, Vaiva Vasiliauskaitė, Julian Hengsteler, Gino Cathomen, Tobias Ruff, Cedric Schmid, János Vörös, Stephan J. Ihle

## Abstract

Systematically perturbing neuronal circuits under controlled conditions offers a promising route to understanding how they encode and process information, yet remains methodologically demanding. *In vitro* neuronal networks can be cultured on microelectrode arrays (MEAs) that allow stimulation and recording, and microfluidic patterning can constrain network topology to yield stable stimulation-evoked responses. Yet, the space of possible spatiotemporal stimulation patterns remains too large for exhaustive exploration. Additionally, the evoked responses depend on prior stimulation history. Here, we present an open-source framework that sustains closed-loop reinforcement-learning (RL) control of engineered neuronal networks, sending electrical stimuli to the MEA and evaluating the evoked responses to identify stimulation patterns that drive specific target activity motifs. We extend inkube, a low-cost, open-source electrophysiology system from off-the-shelf components, with closed-loop electrophysiology. This enables reliable delivery of stimulation at single-sample precision with millisecond-range round-trip times. It also allows independent RL agents to control multiple networks simultaneously. As a proof-of-principle application, we benchmarked different agents on the task of identifying stimulation patterns that maximise the length of clockwise-circular firing sequences. We first demonstrated that stimulation-evoked responses in engineered recurrent networks were stable and separable across the action space over hours of continuous operation. We characterised the dependency of responses on prior stimulation history, finding state dependence in a subset of stimulus pairs. The agents learned to improve reward during training with respect to random stimulation. Agents converged on non-trivial stimulation patterns that span the full action space rather than mirroring the target motif. All hardware designs and software are publicly available, providing an accessible platform for goal-directed functional characterisation of engineered neuronal networks at single-spike resolution.

## Introduction

Understanding how neuronal circuits process information remains a central challenge in neuroscience (1). Even individual biological neurons exhibit computational complexity far exceeding simple somatic summation, with non-linear dendritic processing enabling single cells to classify linearly inseparable inputs (2, 3). How these primitives combine with network topology and recurrent dynamics (4, 5) to produce robust input-output transformations is poorly characterised. Systematic perturbation under controlled conditions, a common approach for studying complex dynamical systems (6), poses several challenges *in vivo*: Complete control over circuit inputs, exhaustive recording of contributing units, and selective reproducible intervention remain difficult to achieve simultaneously (7, 8).

*In vitro* neuronal networks derived from dissociated rat cortical tissue or human induced pluripotent stem cell (iPSC)derived neurons self-organise into functional circuits when cultured on microelectrode arrays (MEAs) (9, 10). MEAs enable non-invasive extracellular recording of action potentials from tens (11) to thousands of electrodes simultaneously (12), while the same electrodes can deliver current or voltage pulses to stimulate subpopulations of neurons. This bidirectional interface provides an experimental platform for studying neuronal computation and plasticity over days or weeks in a controlled setup (13, 14). The network structure can be further constrained through compartmentalisation using polydimethylsiloxane (PDMS) microstructures (15), yielding physically separated populations with constrained connectivity. This facilitates access, increases interpretability of the input-output mapping, and enables systematic characterisation of stable stimulus-response relationships (16, 17). Furthermore, such networks allow for simple computational mappings such as Boolean logic operations (18). However, even in these constrained networks, the space of possible spatiotemporal stimulation patterns grows combinatorially with electrode count and temporal resolution. Furthermore, neuronal behaviour is influenced by the network’s recent spiking history, introducing state-dependent dynamics (19). Consequently, exhaustive exploration of the action space remains intractable.

Previous work has embedded *in vitro* networks in a closedloop setup, sending sensory inputs with a stationary encoding to the culture, decoding its activity as motor outputs, and providing a ‘learning signal’ (20, 21). Several studies have reported improved performance of control tasks when stimulation feedback was used to penalise undesired behaviour. Early work demonstrated that repeated low-frequency stimulation halted upon detection of a predefined response, can yield selective improvements in stimulus±response associations (22), while others used patterned stimulation as aversive feedback, hypothesising that network reconfiguration acts to avoid stimulation (23, 24). More recent approaches have employed various feedback strategies: pre-defined spatial patterns to train pattern recognition (25), prediction-error signals motivated by the free energy principle (26, 27), *in silico* optimised stimulation applied to cortical organoids (28), and optical stimulation pulses to drive homeostatic plasticity (29). However, these paradigms face several open challenges. Long-term retention of learned behaviour has not been reliably demonstrated, and inducing repeatable plasticity effects through electrical stimulation remains elusive across preparations (30, 31), with the underlying mechanisms poorly understood (32). Whether the observed task adaptation constitutes genuine substrate-specific learning or arises from the closedloop paradigm itself remains contested (33, 34).

Several MEA-based electrophysiology setups realised closed-loop stimulation, with open-source (35, 36) or commercially available solutions (37, 38). Implantable neurostimulation devices further exploit closed-loop paradigms for therapeutic intervention (39, 40). Key limitations across these platforms include variable stimulation timing, limited access to internal recording and stimulation communication, and proprietary hardware. A sub-millisecond system for HD-MEAs shows impressive round-trip times, but lacks flexibility and code availability, as it runs directly on a field-programmable gate array (FPGA), which is challenging to synthesise (41). Hitherto, no system combines open-source hardware, deterministic timing of stimulus execution, and easily accessible data streams that facilitate interfacing with complex stimulation algorithms.

Closed-loop control of the activity of biological neuronal networks further requires strategies that cope with non-linearity and non-stationarity without explicit system models. Model-free reinforcement learning (RL) provides a machine-learning (ML) framework, where agents learn optimal policies directly from interaction (42).

RL-based neurostimulation has been explored clinically for vagus nerve (43) and deep brain stimulation (39, 44) and *in silico* in simulated neural systems (45). For *in vitro* networks, tabular and deep RL have been applied to control the number of spikes in a burst on MEAs (46, 47), and adaptive delayed feedback control has been used to disrupt pathological oscillations (48). Activity-dependent adaptation of stimulation patterns with an RL-inspired paradigm yielded selective trialto-trial improvements in predefined stimulus-response associations (22). In all cases, however, control actions operated on timescales of one second or longer, and network readouts were condensed into scalar quantities such as total spike counts, burst rates, or single-electrode response ratios before being passed to the controller, collapsing the spatiotemporal pattern of individual spikes across electrodes.

In this work, we establish an open-source closed-loop electrophysiology platform that interfaces topologically constrained biological neuronal networks with reinforcement learning agents. The system enables deterministic stimulation at single-sample resolution, meaning reliable delivery of the stimulation pulse at exact timepoints, with round-trip times in the millisecond range. Using the inkube platform (49, 50), it simultaneously ensures physiological culturing conditions for day-long experiments. The included software allows for independent agents to control multiple networks simultaneously using RL frameworks. As a proof-of-principle, we define the goal of identifying the stimulation pattern that induces the longest clockwise-circular firing sequence in engineered recurrent neuronal networks. We demonstrate separability and stability of responses and conclude with successful identification of desirable, non-trivial stimulation patterns by various agents. All hardware designs and software are made publicly available.

## Methods

### Cell culturing

#### Chip preparation

Glass-based passive MEAs (60MEA500/30iR-Ti-gr, Multi Channel Systems MCS GmbH, Reutlingen, Germany) were prepared with polydimethylsiloxane (PDMS) microstructures as described previously (16). The two-layered PDMS microstructures, manufactured by Wunderlichips (Zurich, Switzerland) using soft lithography, consisted of open-top nodes for cell bodies connected by microchannels with 4 µm height. The microstructures used 4-node recurrent designs, with one MEA electrode aligned to each microchannel (see Fig. 1a). On one 60-electrode MEA, 15 full networks could be formed, as shown in Fig. 1b. Chip surfaces were coated with poly-D-lysine (PDL), and PDMS membranes were cut from the wafer and aligned to the MEA electrodes using tweezers. The MEA well was filled with phosphate-buffered saline (PBS) and desiccated to remove air bubbles, then PBS was replaced with culture medium before cell seeding.

**Fig. 1.**
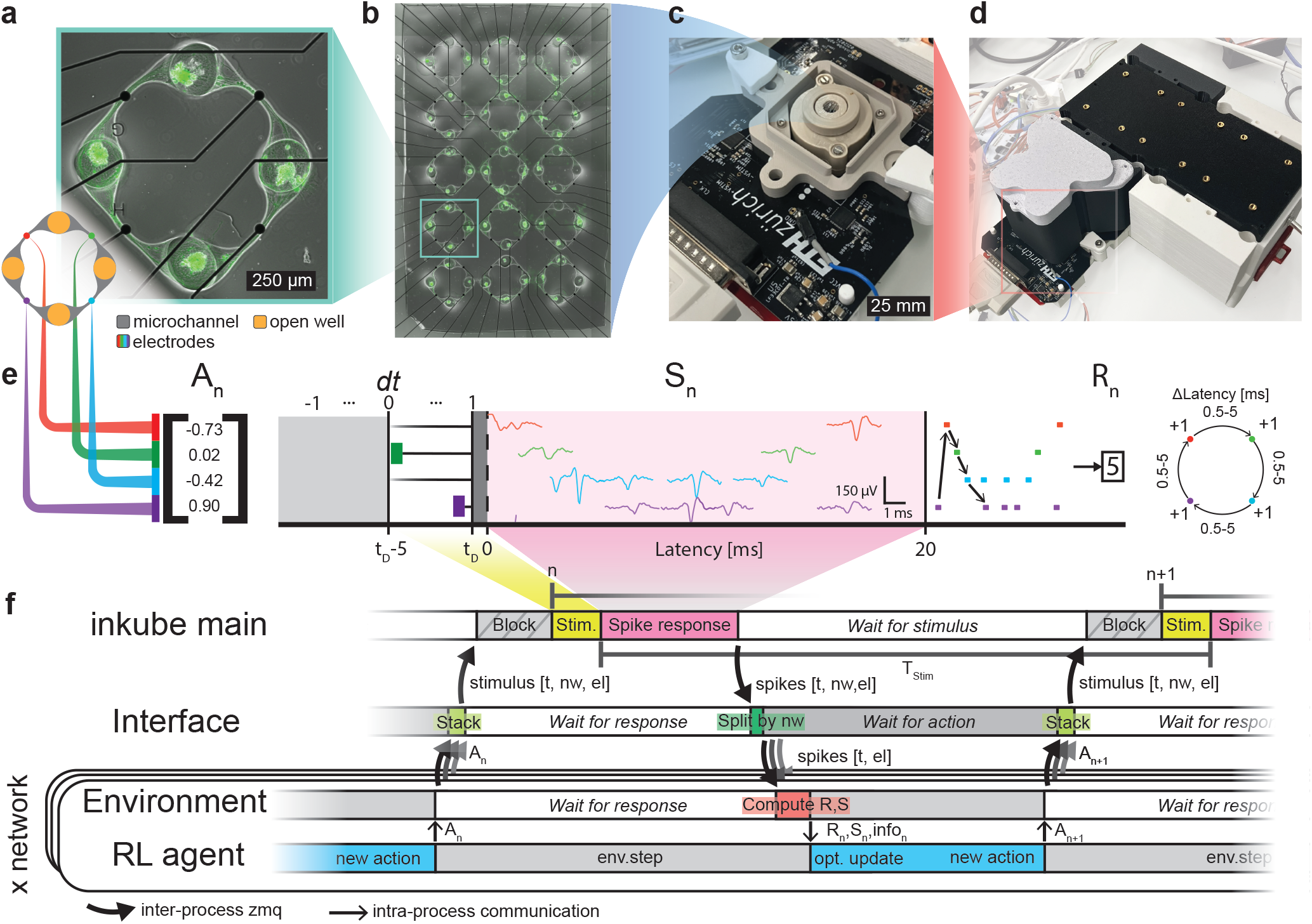
Investigation of *in vitro* patterned networks using a real-time closed-loop stimulation framework with RL control. **a**, *In vitro* neuronal networks in PDMS microstructures on MEAs. Bidirectional circular microstructures confine cell bodies in seeding wells (open at the top) and guide axons through microchannels. Microstructures are aligned such that each microchannel covers one electrode. **b**, 15 networks fit on one 60-electrode MEA. Confocal laser-scanning microscopy image of GFP-expressing Ngn2-induced neurons on DIV 14 that were seeded as spheroids. **c**, Electrophysiology board of inkube. MEA electrodes are recorded via 4 ASICs. **d**, The inkudock system maintains physiological culture conditions. **e**, RL environment implementation on biological networks. A stimulation action A_*n*_ is encoded on each of the 4 electrodes at each time step *n*. Action vector values in [−1, 0) convey no stimulation, whilst values between 0 and 1 are translated linearly into stimulation delays from 5 to 0 ms. A blanking delay *t*_*D*_ is added to all stimulations. Spikes are recorded for 20 ms post-stimulation and constitute the state S_*n*_. For the real-time spike detection, only the cut-out waveforms are stored. The waveforms are plotted in the subfigure. The reward R_*n*_ is the length of the longest valid clockwise sequence found across all possible starting electrodes, where consecutive spikes on successive electrodes must fall within a 0.5-5 ms inter-spike interval corresponding to the expected synaptic transmission window. **f**, Timing of inkube, interface, and RL process communication. The RL process consists of the environment for inter-process communication and the RL agent. The agent sends an action vector for its respective network for the period *n* through the step command. It is forwarded via the environment to the interface, which translates and stacks the actions from all agents into a *m* × 3 matrix, with *m* being the number of stimuli and the columns corresponding to timing, target network, and electrode. Once the interface has received an action from all RL processes or approaches the block time, the stimulus vector is sent out. The inkube main process applies the stimulus, given that it is received before a block phase for internal communication. After recording, the spikes are split by network and distributed to the environments, which locally compute the reward and state based on their settings. The agent can incorporate the feedback information into its action choice by updating its value functions. This cycle repeats at a frequency 1*/T*_Stim_.

#### Culture preparation

Early-stage Ngn2-induced neurons derived from human induced pluripotent stem cells (iPSCs, provided by Novartis, Basel, Switzerland) were thawed and differentiated in neurobasal differentiation medium (NBD+) following established protocols (50, 51). Two seeding strategies were used. For spheroid seeding, 100 cells per spheroid were aggregated using AggreWell^*TM*^ 400 plates (34415, StemCell Technologies, Vancouver, Canada) and plated individually into the microstructure wells on the MEA after 24 h according to the manufacturer’s instructions. For suspension seeding, dissociated cells were plated directly at a density of 300,000 cells per MEA. In both cases, medium was supplemented with 2 µg/mL doxycycline (631311, Takara Bio USA, Inc., Mountain View, CA, USA) and laminin (L2020, Sigma Aldrich, St. Louis, MO, USA) for the first week. Culture medium was exchanged twice a week by replacing half of the medium with fresh NBD+.

### Experimental setup

#### Custom electrophysiology

Stimulation and recording of the MEA electrodes were achieved with the Intan RHS2116 ICs (Intan Technologies, Los Angeles CA, USA), controlled by a system-on-chip (SoC) (XC7Z020-1CLG400C, Xilinx, San Jose, CA, USA). We used these components in a further optimised modular version of the inkube system, which we presented in earlier work (49). Up to 4 recording boards housing 4 of the electrophysiology chips, each interfacing one 60-electrode MEA, can be attached to the SoC (see Fig. 1c and Fig. S3). The sampling frequency was set to *f*_S_ = 17, 361 Hz. Digital low-voltage differential signalling (LVDS) communication and supply voltages were connected through a 25-pin DSUB from the SoC shield. Positive and negative stimulation voltages were generated on the power board, the supply voltage was generated on the electrophysiology board with a low-dropout regulator (LDO) from 5 V supplied by the SoC.

#### Custom incubator

All experiments were performed with ‘inkudock’, a custom incubator with a central reservoir and up to 4 attached chambers, described in previous work (50) and shown in Fig. 1d. An updated version of the recording unit holder was employed to accommodate the glass-based MEAs (for design files and assembly see Supplementary Information B.1 and Fig. S2). A temperature sensor was mounted on top of the base printed circuit board (PCB), controlling the Peltier thermoelectric device located underneath. The electrophysiology boards were mounted on top of the MEA using spring contacts (Supplementary Information B.5 and Fig. S4a-c). A passive chamber enclosed the culture and connected it to the reservoir (Supplementary Information B.3 and Fig. S4d-f). A lid with a water compartment on top of the culture prevented evaporation in the non-humidified environment (50). Following a calibration test, a medium temperature of *≈*35.5 °C was achieved by setting the MEA holder temperature to 32.3 °C at a reservoir temperature of 42 °C (for details see Supplementary Information B.6 and Fig. S5).

### Closed-loop electrophysiology

#### Spike detection

Raw voltage traces were acquired from UDP packets sent by the SoC at *f*_*S*_ and processed by a C++ pipeline (see Supplementary Information A.2 and Fig. S1). Signals were high-pass filtered (300 Hz cutoff, 2nd-order Butterworth) with spike detection performed using an adaptive threshold set to 6*×* the median absolute deviation of the filtered signal, updated at 1 Hz with exponential moving average smoothing (*α* = 0.25). Upon negative threshold crossing, the local minimum was identified within 20 samples, followed by a blinding period of about 1.5 ms. Detected spikes were transmitted to the Python control program (`inkube_main`) containing channel ID, timing, and a 45-sample waveform (2.6 ms) (see Supplementary Information A.5).

#### Stimulation and artefact suppression

Stimulation pulses were charge-balanced biphasic waveforms (positive-first, 461 µs per phase, 50 nA amplitude). Pulses reliably induced a network response (for post-stimulation time histograms at varying amplitudes see Fig. S7). Hardware artefact suppression combined active discharge (*∼*1.5 ms post-pulse), highpass filter switching (increased to 1 kHz during stimulation), and discharging amplifier outputs via fast-settling. The stimulation offset *t*_*D*_ was set to 2.1 ms to accommodate pulse execution and discharge. Spike detection threshold updates were suspended during stimulation periods to prevent artefact-driven threshold adaptation.

#### Communication and segmentation

Closed-loop stimulation was achieved by segmenting experiment time according to the desired stimulation frequency (Fig. 1f). A 20 ms post-stimulus window collected incoming spikes, which were transmitted to the `RL_Interface` program via ZeroMQ sockets (52) (see Supplementary Information A.3). The `RL_Interface` aggregated activity by network, enabling individual reinforcement learning agents to query responses and return actions. Actions were translated into stimulation parameters (electrode selection, timing delays, amplitudes) and transmitted to `inkube_main` at period end or when all agents responded. Commands were encoded as register-write operations for the Intan chips and sent via USB 2.0 bulk transfers (see Supplementary Information A.4).

#### Reinforcement learning environment

To derive stimulation strategies that elicit desired spiking responses, we formulated the task as a reinforcement learning (RL) problem. In RL, an agent learns a policy that maximises cumulative reward through interaction with an environment. Here, the biological neuronal network serves as the environment, whilst a Python program acts as the agent selecting stimulation parameters. The reward function, defined on the recorded spiking activity, can be freely designed to implement arbitrary goals.

The experimental protocol proceeds in discrete cycles (Fig. 1e-f): The agent selects an action in the form of an electrical stimulus, the network’s induced response in spiking activity is recorded for 20 ms, this response is transformed into a state representation, and a reward is computed. Based on this feedback, the agent selects the next action, and the cycle repeats at the set stimulation frequency.

Mathematically, we model this as a Markov decision process (MDP) defined by the tuple (*S, A, R, T* ), where *S* is the state space, *A* is the action space, *R* is the reward function, and *T* represents the transition dynamics of the biological network. The reward function *R* is known and depends only on the network state, not the applied action. The transition dynamics *T*, however, are unknown and must be learned through interaction.

This formalism makes several assumptions that likely do not strictly hold for biological networks. First, we assume the network is stationary, though synaptic plasticity and other long-term changes may occur. Second, we discretise time into observation windows, though neuronal dynamics are continuous. Third, we define the state operationally as the post-stimulus spiking response recorded for 20 ms at the four microchannel electrodes. This captures only a projection of the underlying network dynamics. Sub-threshold membrane potentials, chemical signalling, and activity of neurons not captured by the electrodes are not represented. The MDP framework assumes that the state is sufficient to predict future transitions (the Markov property), which is unlikely to strictly hold for this reduced representation. In practice, we treat the system as an MDP and rely on the RL agents to learn useful policies from the available observations. Despite these simplifications, the MDP framework provides a tractable approach to learning control policies for biological neuronal networks. The following sections provide a detailed description of the required elements and our design choices.

#### Reward computation

Reward computation defines a measure of how desirable a given behaviour is at each time step; here, it depends solely on the state, not the action taken. Rewards must be designed to maximise desired agent behaviour through dense, non-sparse feedback that eases credit assignment (53).

Unless otherwise stated, a reward was used that promotes clockwise firing. For each electrode present in the response, a sequence search from its earliest spike was performed by iteratively checking for valid continuation spikes on the next electrode in clockwise order. A spike was considered a valid continuation if it occurred within a window of 0.5 ms and 5.0 ms, as this is the time frame expected for synaptic transmission and conduction to the next well. When multiple spikes satisfied these timing constraints, the earliest valid spike was selected. The reward was defined as the maximum sequence length obtained across all possible starting electrodes. This approach prioritised sustained circular propagation whilst remaining robust to spontaneous activity.

#### Action space

The action space in RL can either be discrete or continuous. Fundamentally different methods are required to train both variants. In this work, both discrete and continuous action spaces have been investigated.

The action vector was a four-dimensional vector in [*−*1, 1]^4^, where each element encoded the timing of an electrical stimulus on one of four network electrodes, as shown in Fig. 1e. Negative numbers encoded the lack of a stimulus, while positive values encoded the stimulus timing relative to recording start: The action approaching 0 from the positive side corresponds to maximum latency (5 ms) and 1 to minimum latency (0 ms), with an additional fixed delay of *t*_*D*_ applied to all stimuli, which corresponded to the duration of one stimulation pulse. Given the system’s sampling frequency, actions could be applied with a single sample timing precision of approximately 58 µs.

For discrete action spaces, *n*_action_ values were uniformly sampled per electrode at intervals of 1*/*(*n*_action_ *−* 1), always including 0 and 1. In this work, we set *n*_action_ = 4, yielding a total action space of (*n*_action_ + 1)^4^ = 625 possible action combinations, where 4 is the number of electrodes. All randomly generated action vectors for system benchmarking contained at least one non-negative entry.

#### State space

The post-stimulus response was recorded for *T*_S_ = 20 ms and the spikes on the 4 electrodes were captured, defining the network state S_n_. The full state was therefore represented by a sparse, binary matrix of size *n*_electrodes_ *×*(*T*_S_ *· f*_S_).

RL is generally data-intensive to train (5). In our case, data collection is constrained by the physical experimental setup, which imposes limits on the maximum stimulation frequency and thus the rate of sample acquisition. Therefore, agent complexity was reduced by restricting the state dimensionality using two different compression methods that condensed the spiking dynamics to a continuous latent space of dimension *d*_state_. We implemented a linear encoding with PCA and a deep learning based non-linear encoding with a convolutional neural network. The state encodings were fitted on 7200 samples generated by Latin hypercube sampling (54) across the action space (continuous or discretised). The sampling targets adequate coverage of possible network responses and was performed at least every 24 h to account for drift in network behaviour.

#### Principal-component analysis (PCA)

For PCA, spike time binning at three different temporal resolutions was tested: 0.29, 1, and 2 ms. The binning produced a binary feature vector of length 4 *× n*_bins_, where each element indicated the presence of a spike on a given electrode within the *n*_bins_ time bins. Principal component analysis was applied to reduce this to the target state dimensionality.

#### Dilated 1D convolutional neural network (DCNN)

A lightweight temporal convolutional network was trained to encode spike sequences into low-dimensional state representations (55). The network processed up to 30 spikes, where each spike was represented by a one-hot encoding of its electrode identity (4 dimensions) and the square root of its spike time (1 dimension). The architecture employed depthwise separable convolutions and dilated kernels to capture multi-scale temporal patterns whilst maintaining 97 trainable parameters. The network was trained using a pairwise repulsion loss that maximised distances between state representations of different spike patterns, encouraging distinct encodings. Training continued for up to 200 epochs with early stopping when the mean pairwise distance exceeded a threshold of *−*1.0.

#### Gymnasium environment

In order to facilitate agent testing as well as interaction with the cultures, two custom Gymnasium environments for continuous and discrete action spaces were designed, requiring definition of the poststimulus recording length, reward function, and state representation (56). Each environment controlled a single network on an MEA as shown in Fig. 1f.

The environments communicated with the `RL_Interface` program via ZeroMQ sockets. Each `env.step(action)` call queued an action for the next available stimulation slot within the segmented time structure described above. The environment returned the standard Gymnasium tuple (state, reward, terminated, truncated, info) once the post-stimulus window completed and spikes were aggregated. Actions were validated and formatted as stimulation parameters (timing and location) by the `RL_Interface` before transmission. Invalid actions defaulted to the null action (no stimulation) but still returned a valid tuple to prevent pipeline interruption. Example usage is shown in the Supplementary Information A.6.

The `info` dictionary provided diagnostic data, including spike times in a sparse encoding, missed stimulation events since the last call, and pipeline status flags. A minimum of 8 ms for agent computation and inter-process communication was allocated at 20 Hz, increasing to 208 ms at 4 Hz. State representations requiring computationally expensive operations therefore required lower stimulation frequencies. The returned action field allowed verification that the intended stimulus was applied.

#### RL agents

As we use both discrete and continuous action spaces, which require fundamentally different training methods, we developed different agents for the two types. Discrete agents selected from the 625 predefined action combinations described above, while continuous agents operated over the full continuous space (only limited by finite numerical precision), which were mapped onto the discrete delay by the interface (*≈* 60, 000, 000 combinations).

We compared agents using full state information (state-based) with those relying only on action history (state-free). As training steps required for convergence varied drastically, values ranged from 10k to 120k, always followed by 4800 (20 min) steps of performance evaluation.

#### Random agent

The random agent acts as a control, uniformly sampling the continuous action space. Due to the action encoding, it stimulates each electrode with a probability of 50 %.

#### Multi-armed bandit (MAB) agent

The multi-armed bandit (MAB) is a state-free RL formulation in which the agent maintains an estimate of the expected reward for each action and balances exploration of untested actions against exploitation of the current best.

The MAB agent operates on the discretised action space described above and selects an action *a*_*t*_ using the upper confidence bound (UCB):

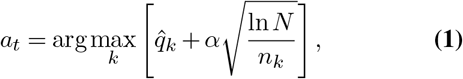

where *k* identifies a unique action with 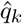being the empirically estimated expected reward of this action, *N* the total number of actions applied, *n*_*k*_ the number of times action *k* has been selected, and *α* a scaling factor controlling the degree of exploration. For this work, *α* = 0.5.

To extend the MAB to continuous action spaces, we implemented an adaptive variant that initialises with the same 625 discrete arms but periodically refines them using a cross-entropy method (CEM). Every 2,500 steps, the arms are ranked by their estimated reward. A Gaussian distribution is fitted to the 125 highest-performing arms (elite set), and 250 new candidate arms are sampled from this distribution, replacing the lowest-ranked arms whose statistics are reset. Actions are clipped to the valid range [*−*1, 1]^4^. This procedure progressively shifts the arm population from the initial discrete grid towards high-reward regions of the continuous action space.

#### Linear contextual bandit (LCB) agents

The LCB extends the MAB framework by conditioning action selection on the current network state approximated through the previous action response. Both discrete and continuous variants were implemented.

In the discrete case, the agent operates on the same discretised action space as the standard MAB and estimates a state-dependent value function:

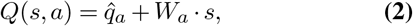

where 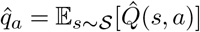 is the empirical expected reward of action *a* as computed in the discrete MAB agent, and *W*_*a*_ is a weight vector capturing the linear dependence of the reward on the state. Intuitively, 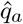 represents the average reward of an action across all states, while *W*_*a*_ *· s* corrects this estimate based on the current network state. *W*_*a*_ is refitted every 100 samples via the least-squares solution:

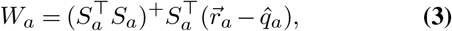

where 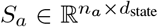 is the matrix of all states observed when action *a* was applied *n*_*a*_ times, (*·*)^+^ denotes the Moore-Penrose pseudoinverse, 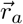 is the vector of all rewards received for action *a* (one entry per trial), and 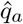 is subtracted so that *W*_*a*_ is fitted to the reward residuals rather than the raw rewards. Exploration follows the same UCB strategy as the discrete MAB with an exploration value *α* = 5. We additionally implemented a two-phase variant, termed Dynamic LCB, in which the first phase trains only 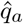; this estimate is then frozen in the second phase, during which only *W*_*a*_ is fitted. This separation prevents early, poorly estimated state corrections from corrupting the action-value baseline.

In the continuous case, Equation 2 cannot be used directly: because *Q* is linear in *a*, optimisation would always drive the action to the boundary of the action space rather than finding an interior optimum. To allow the model to capture optima within the action space, the reward is instead expressed as a linear function of an expanded feature vector that includes nonlinear terms:

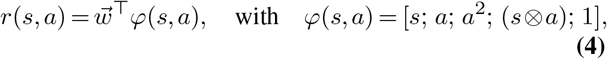

where ⊗ denotes the outer product. The quadratic term *a*^2^ introduces curvature so that the reward surface can peak at intermediate action values, while the interaction term *s* ⊗ *a* allows the optimal action to shift depending on the current network state. The bias term 1 captures any constant offset. Although the model is linear in the weights 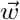, the nonlinear feature expansion allows it to represent a quadratic reward landscape over actions.

The weight vector 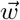 is estimated using Bayesian linear regression, which maintains a Gaussian posterior that is updated incrementally as new observations arrive:

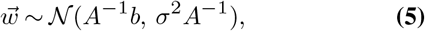

where *A* ∈ ℝ^*p*×*p*^ accumulates the outer products of observed feature vectors and *b* ∈ ℝ^*p*^ accumulates the reward-weighted feature vectors, with *p* the dimensionality of *φ*. Both are initialised as the identity matrix and zero vector, respectively, corresponding to a Gaussian isotropic prior. After each trial with observed feature vector *φ* and reward *r*, the posterior is updated as *A* ← *A*+ *φφ*^⊤^ and *b* ← *b* + *r φ*, incorporating the new data point without requiring access to the full history.

Action selection during exploration follows Thompson Sampling: a weight vector 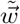 is drawn from the current posterior, and the action maximising 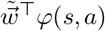 is selected. Because the feature vector contains no cross-electrode terms *a*_*i*_ *· a*_*j*_, this optimisation decomposes into independent per-electrode quadratics that can each be solved in closed form. Sampling from the posterior introduces directed exploration, as uncertain weight estimates produce variable action preferences, naturally favouring under-explored regions of the action space. During exploitation, the posterior mean *A*^−1^*b* is used in place of a sample, selecting the action with the highest expected reward under the current model.

### Data analysis

#### State dependency

To assess the statistical significance of state-dependent modulation, we tested whether the compressed current states (**Ŝ**_*n*_) depended on both the current action (**A**_*n*_) and the previous action (**A**_*n*−1_). For each network and action pair, we partitioned **Ŝ**_*n*_ into four groups based on the combinations of current and previous actions. We then computed the distributions of each state dimension within these groups and quantified dissimilarity between the multidimensional distributions sharing the same current action **A**_*n*_ but differing in previous action **A**_*n*−1_ using the Sliced Wasserstein Distance (SWD) metric. The SWD was computed by projecting both *d*_state_-dimensional state clouds onto *P* = 200 random unit vectors sampled uniformly from the (*d*_state_ *−* 1)-sphere, computing the exact one-dimensional Wasserstein distance along each projection, and averaging across projections. This yields a single scalar dissimilarity measure per action pair that is sensitive to differences in any linear direction of the state space, without requiring multiple comparisons across dimensions.

To establish statistical significance, we generated null distributions through temporal scrambling of the previous action identity (**A**_*n*−1_) whilst maintaining the current action (**A**_*n*_) and neural response pairings intact. For each network-action pair, we computed the SWD for 200 random permutations of **A**_*n*−1_ to establish a null distribution under the null hypothesis of no state dependency. Z-scores were calculated by comparing the Wasserstein distance of **A**_*n*−1_ against the mean and standard deviation of the scrambled distribution. Network and action pairs exceeding the significance threshold (*z >* 2.326 for *p <* 0.01) were considered to exhibit statistically significant state-dependent effects. The proportion of action pairs reaching significance was tested against the expected 1 % false positive rate using a one-sided binomial test with Benjamini±Hochberg correction applied across circuits.

#### Agent comparison

To compare agent performance, only networks with a mean electrode firing rate exceeding 4 Hz during spontaneous recording were included. Two metrics were computed from the test phase: mean test reward and top 20% test reward, defined as the mean of the highest-scoring 20% of test samples, capturing peak rather than average performance. For statistical comparison, data recorded across multiple days in vitro were averaged per agent±network combination to yield a single representative value, which was then min-max normalised within each network to account for differences in baseline activity between preparations. Pairwise comparisons between agents were performed using the twosided Wilcoxon signed-rank test. For agent pairs evaluated across both experimental batches, all shared networks were included in the paired test. Statistical significance was assessed at *α* = 0.05.

## Results and Discussion

### Open-source platform enables day-long closed-loop electrophysiology

Closed-loop electrophysiology requires adaptation of the applied stimulation based on culture activity at physiological timescales. For this, the full signal processing chain with data acquisition, spike detection, agent computation, and stimulation delivery needs to complete within the inter-stimulus interval. We achieved this by extending the open-source inkube system (49), built around an electrophysiology ASIC controlled by a SoC that communicates with a host PC for interpreting voltage recordings and receiving commands. First, we limited the UDP link to acquisition of data and added USB communication for commands (Fig. S1). Furthermore, we developed a novel software with a spike detection pipeline implemented in C++, a Python-based RL framework with Gymnasium-compatible environments (56), and inter-process communication via ZeroMQ (52) (for implementation details see Methods).

The spike detection pipeline achieved a latency of less than 1.2 ms from raw data acquisition to event publication. Stimulation commands were transmitted to the system before each period such that pulses were executed at single-sample precision (*≈* 58 µs at *f*_*S*_ = 17,361 Hz) independent of communication jitter. Maximum closed-loop stimulation frequency was set to 20 Hz, comprising response collection (20 ms), spike detection (<1.2 ms), block phase reserved for USB communication (*≈* 13 ms), stimulation (5 ms) and offset (t_*D*_), leaving at least 8 ms for inter-process communication and agent computation. The minimal system round-trip time was determined by triggering stimulation upon a detected spike on a designated electrode and is presented in Fig. S6.

Long-term culture viability was ensured by the inkudock incubation system (50), which provided temperature control and evaporation prevention (Fig. 1d). The platform sustained continuous closed-loop operation exceeding 33 h. All six agents were trained and tested sequentially in a single uninterrupted session, comprising 5 min of spontaneous recording, state encoding calibration, and successive training runs followed by test phases, all at 4 Hz. Up to four MEAs were operated simultaneously, each hosting 15 independent networks with individual RL agents, yielding up to 60 concurrent closed-loop experiments.

### Evoked responses show reward stability and separability over hours

Before deploying RL agents, we characterised the range and temporal stability of network responses across the action space. We generated 500 random stimulation patterns (actions) and applied them by drawing uniformly at random at 4 Hz, for a total of 50,000 stimulations per network, corresponding to approximately 3.5 h (Fig. 2a). On average, 125 s (500/4 Hz) separated two occurrences of the same action, and each action was applied about 100 times. Example reward traces for four actions on a single network are shown in Fig. 2b, representing the actions with the highest and lowest mean reward and reward variance. Post-stimulus raster plots (Fig. 2c) reveal several consistent response bands, where network electrodes fire at a characteristic latency after stimulation. Despite hundreds of intervening stimuli between repetitions, these spatiotemporal response profiles show stable relative timing of spikes across occurrences of the same stimulus. We can therefore assume that responses are largely driven by the current stimulus. If the influence of the stimulation history was substantial, there would be significant variations in the response to each individual occurrence of a specific action. No systematic drift during early repetitions is apparent, consistent with prior observations in similar systems at the same stimulation frequency (16, 17). Post-stimulation histograms commonly show differences in jitter between directly induced axonal and synaptically transmitted activity (57). As the stimulation window spans up to 5 ms with added blanking, early responses were a mix between directly induced spikes and synaptically transmitted activity, while late responses were less reliably induced when averaged across occurrences (see example histograms in Fig. S8and analysis in Fig. S9).

**Fig. 2.**
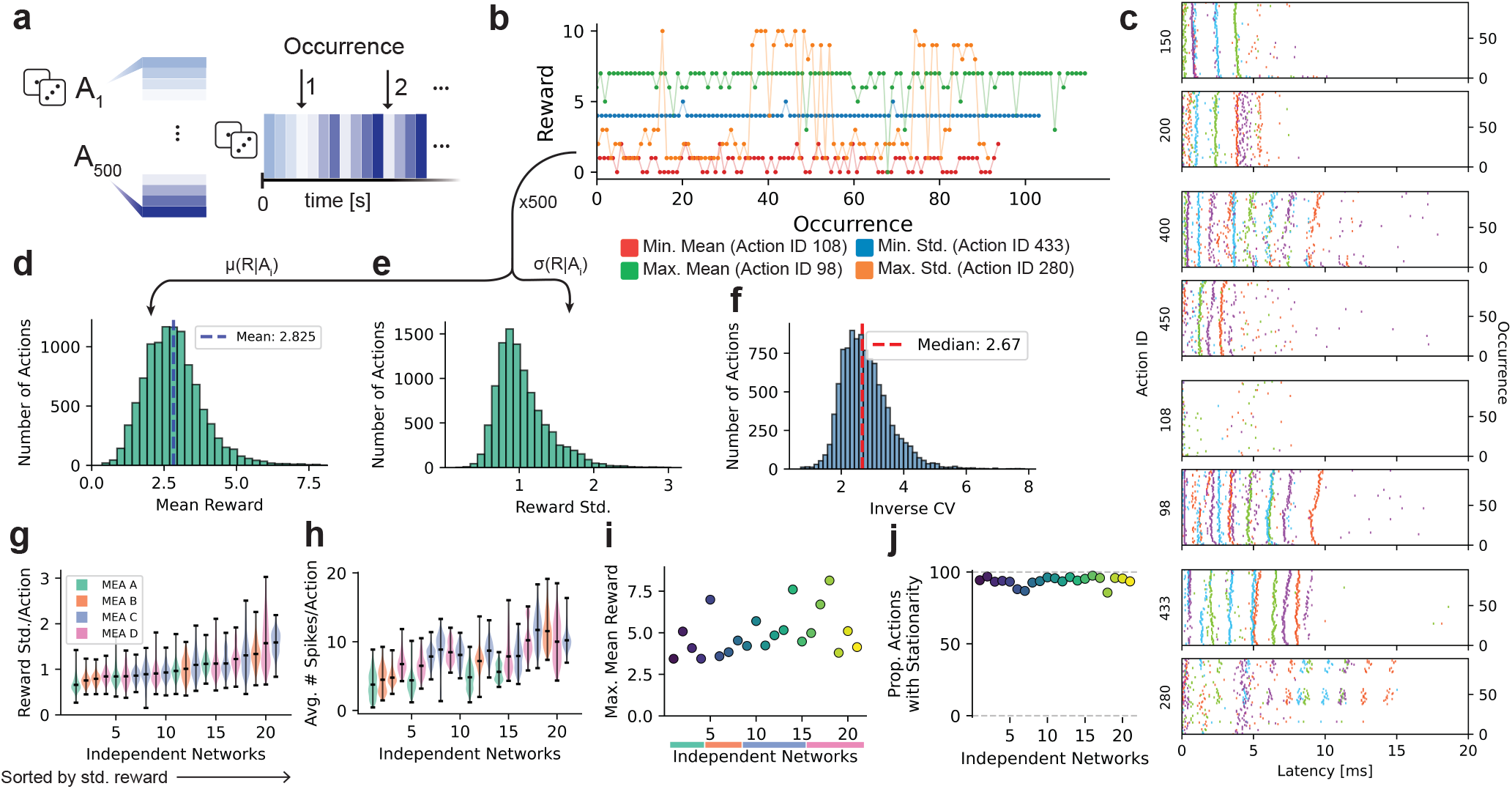
Responses are stable and separable across the action space. **a**, 500 random action vectors were generated, with at least one electrode stimulated. Actions were drawn uniformly at random and applied at 4 Hz for a total of 50,000 stimulations per network (each action was applied around 100 times). **b**, The reward associated with the same action was extracted across repetitions. The four shown actions represent the mean reward and reward standard deviation maxima and minima of a single network on DIV 24. **c**, Post-stimulus raster plots are shown for selected actions and the extrema actions from **b** across repetitions on the same network as in **b**. A spike on one of the network electrodes after stimulation is marked by a dot at its latency, colour-coded by electrode as in Fig. 1a. **d**, Action performance was compared for all actions on all networks, averaged across repetitions. The distribution of mean rewards peaks near 2.8, meaning the mode of the distribution corresponds to more than half a cycle inside the microstructure. **e**, The reward standard deviation was calculated for all actions and all networks across repetitions, showing a right-skewed distribution centred around 0.9. **f**, The inverse coefficient of variation (mean reward divided by standard deviation) was computed for all actions and networks. The median of 2.67 indicates that for most actions, the mean reward exceeds the trial-to-trial variability by more than twofold. **g**, Reward standard deviation was summarised per network across four MEAs (MEA A spheroid seeding, B-D suspension seeding). In total, we included 21 networks in our analysis, which showed an average firing rate on all 4 electrodes of at least 4 Hz while recording spontaneous activity and good signal quality. Independent networks were sorted by median standard deviation, showing significant differences in variability between networks. **h**, The mean number of evoked spikes per action per network was analysed. Networks with higher spike counts tend to show higher reward variability. **i**, To assess the range of achievable performance, the maximum mean reward across all 500 actions is shown for each of the networks. Networks are now sorted by MEA as indicated by the colour code. This encoding is kept consistent with Fig. 4a,c. **j**, To verify that reward signals remain stable over the full duration of the experiment, the Augmented Dickey-Fuller test for stationarity was applied to the reward sequence of each action. Across networks, approximately 90 % of actions show stationary responses (*p <* 0.01), confirming that the evoked responses do not systematically drift over the course of the experiment.

Across all actions and networks, mean rewards span from 0.5 to approximately 8, with the distribution peaking at around 2.8 (Fig. 2d). The reward standard deviation across repetitions is right-skewed, centred around 0.9 (Fig. 2e). To assess whether actions are sufficiently separable for RL agent training, we computed the inverse coefficient of variation (mean divided by the standard deviation of the reward) for each action (Fig. 2f). The distribution has a median of 2.67, indicating that for most actions the reward signal exceeds the trial-to-trial variability by more than twofold. This suggests that a reliable estimate of the expected reward for an action, the action value, can be learned by the RL agents after only a few repetitions of the same action. Ranking the actions relative to each other early on is therefore possible (for action discriminability see Fig. S20). As the reward metric evaluates spike sequences within defined temporal windows, it is robust against spontaneous activity occurring outside these windows and on other electrodes. The distribution from the control without cells is shown in the Supplementary Information C.6, showing a peak in the reward at 1 and a normally distributed standard deviation with a mean around 0.5.

Despite sharing the same microstructure design, independent networks differ in their evoked spike counts per action (Fig. 2h). A higher expected spike count correlates with higher reward variability (see Fig. S12 for correlation analysis). Trial-to-trial variability in evoked responses is expected from intrinsic excitability fluctuations at the single-neuron level (58) and synaptic dynamics in network-embedded neurons (19). A higher number of synaptic connections may lead to more probabilistic contributions to the reward. The maximum achievable mean reward also varies across networks (Fig. 2i). Potential underlying causes for inter-network differences include varying cell counts, the proportion of stimulated units, or differences in network synchronisation. To verify that reward signals remain stable over the full duration of the experiment, we performed the Augmented Dickey-Fuller test for stationarity on the reward sequence of each action.

Across all networks, approximately 90 % of actions showed stationary rewards (*p <* 0.01) in Fig. 2j. This does not imply stationarity of the network itself. However, for a high proportion of the randomly sampled actions, any potential network non-stationarity does not affect the reward metric.

To test the impact of residual stimulation artefacts, we performed the same protocol on an MEA with a microstructure without cells in cell culture medium. Representative raster plots are shown in Fig. S13with the full post-stimulation histogram in Fig. S14. Artefacts have consistent timing across occurrences. Many raster plots have a visible stimulation artefact below 0.5 ms, which is likely caused by the filter ripple resulting from the switch-off of the active discharging. In some cases, additional bands can be seen around 1 to 2 ms, which is potentially due to residual charge decay causing a ripple in the IIR-filter response. The distribution of associated average rewards and standard deviation is shown in Fig. S15. The standard deviation is much lower than with cells, and maximum rewards reach up to 3, caused by the low minimum window time of the reward metric.

### Action responses for selected actions exhibit dependency on the previous action

Recruitment of axons through stimulation depends on previous activity (58). As also visible in the example raster plots in Fig. 2c, some synaptic connections in the response appear or disappear over stimulus iterations. We therefore assessed whether the response to a stimulus is influenced by the spatiotemporal pattern of the previously applied stimulus. Pairs of random stimuli *X* and *Y* were generated. These were applied in a random order, and the responses *S*_*n*_ were compressed using different state compression algorithms. For each pair, 1920 stimuli were applied. The responses were summarised into groups with the same current and previous action. The distributions with the same current action were analysed for their difference induced by the previous action by comparing them to scrambled distributions (Fig. 3a-c, details provided in Methods). Pairs with z-scores exceeding the value corresponding to the 99th percentile were considered significant.

**Fig. 3.**
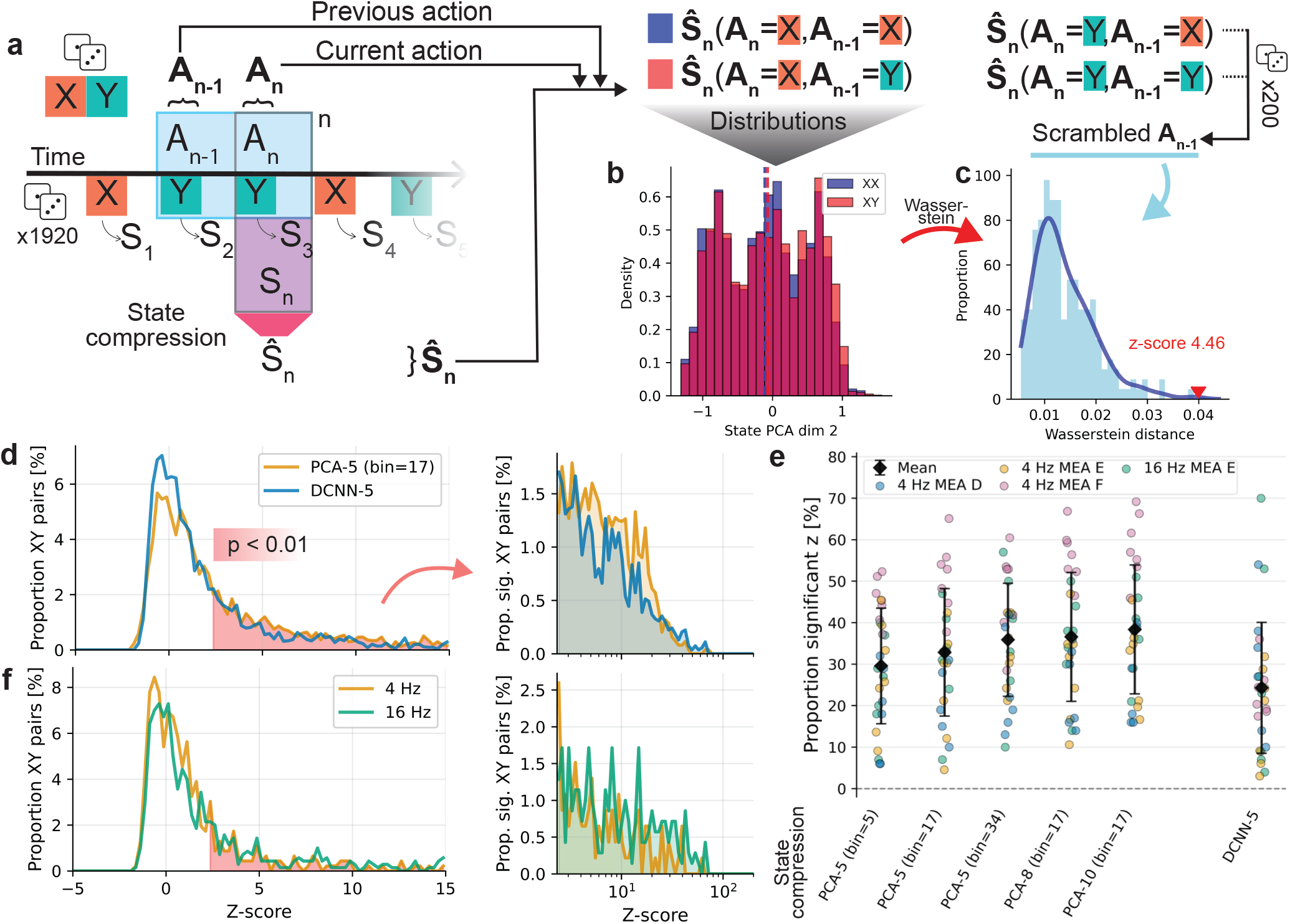
State representation of responses is separable by the stimulation history. **a**, A pair of random actions X and Y was generated and applied in a random order to test whether the previous action can be determined from the network response after the current action. Action pairs that stimulate at least one electrode were generated and applied 1920 times. The post-stimulus response S_n_ as defined in Fig. 1e was recorded and compressed to Ŝ_n_ using a state compression algorithm. With the state we stored the current action A_n_ and previous action A_n−1_ . The elements were summarised into vectors **Ŝ**_**n**_, **A**_**n**_, and **A**_**n**−**1**_ for each action pair. **Ŝ**_**n**_ was then split into 4 groups by the different current and previous actions: XX, XY, YX, YY. **b**, The distributions for every state dimension of each subset of **Ŝ**_**n**_ were calculated. Subsequently, the distributions of the states with the same current action **A**_**n**_ were compared in the *n*_state_-dimensional space using the Sliced Wasserstein distance metric (see Methods for details). **c**, The distances of the subsets split by previous action **A**_**n**−**1**_ were compared with a null distribution, obtained by arbitrarily splitting with a scrambled vector **A**_**n**−**1**_ . The z-score of each action with respect to the scrambled distribution was extracted. Based on this, we analysed whether the previous action led to a larger distance of the distributions of a group when split by the previous action instead of a scrambled vector. A larger distance between the groups when split by previous action means discriminability, and therefore a dependence of the state on the previous action. **d**, The z-score of each pair of actions in comparison with its respective null distribution was tested for significance (p < 0.01; *z >* 2.326). The significant proportion is shown on a logarithmic scale. The PCA approach has 35 % of significant z-scores, the DCNN approach 26.2 %, meaning those proportions of the tested action pairs enabled separation of the state distributions based on the previous action. **e**, The proportion of significant action pairs was compared across PCA and DCNN state compression methods with varying dimensionality and binning parameters. The data was aggregated by network and stimulation frequency (6 networks on MEA D, random seeding; 7 networks each on MEA E and F, spheroid seeding). The different number of XY pairs tested were 50 (4 Hz; MEA D), 32 (4 Hz; MEA E), 50 (16 Hz; MEA E), and 86 (4Hz; MEA F), for the 4 datasets. **f**, Results obtained at stimulation frequencies of 4 and 16 Hz were compared using DCNN-5. The significant proportion is shown on a logarithmic scale. At 16 Hz, a significantly larger proportion of action pairs yielded separable network states (29.6 % vs. 19.3 %, two-proportion z-test, *p <* 0.001), suggesting that higher stimulation frequency increases state dependence.

The distributions of 218 stimulus pairs at two different stimulation frequencies across different networks on different MEAs are shown in Fig. 3d for two different state compression algorithms. Both show significant differences in the state depending on the previous action for about a third of the pairs. This indicates that both algorithms capture relevant differences that persist across the stimulation period.

The state compression algorithms were compared when averaged by network and stimulation frequency. To assess whether each metric yielded more separable state pairs than expected by chance, a one-sample binomial test was performed against a null proportion of 1 % (corresponding to *p <* 0.01 under the standard normal), with Benjamini-Hochberg correction across metrics. Both methods achieved statistically significant separation (*p <* 0.001). To validate that significant effects reflected genuine lag-1 state dependency rather than longer-range temporal autocorrelation inherent to the network dynamics, we repeated the analysis, substituting *A*_*n*−1_ by *A*_*n*−5_ and *A*_*n*−10_ as control. For the multi-step lag analysis, no significant dependencies were obtained (see Fig. S17).

PCA performance varied slightly across binning parameters. Increasing the number of considered principal components yielded only minor improvements. The temporal convolutional approach (DCNN) showed slightly worse performance, achieving separation on average in more than 26 % of the action pairs averaged across circuits. Increasing the stimulation frequency to 16 Hz showed significantly better separability for DCNN (see distributions in Fig. 3f; two-proportion z-test) relative to 4 Hz stimulation. However, experiments were performed at 4 Hz to avoid affecting the network responses over time, which is expected at 16 Hz stimulation (17). As the DCNN also takes substantial training time, PCA was selected for the experiments.

Single-neuron excitability fluctuates over extended timescales due to intrinsic ion channel dynamics (58), and synaptic dynamics contribute an additional source of trial-to-trial response variability in network-embedded neurons (19). In comparable patterned networks, Duru *et al*. observed only minor frequency-dependent effects at 1±4 Hz, with latency shifts and spike count reductions appearing at 8 Hz and above (17). Our results go beyond frequency-dependent adaptation by showing that even at 4 Hz, the specific spatial pattern of the preceding stimulus can measurably influence the current response in a subset of action pairs. In the microchannels, we expect to recruit a large proportion of the axons during stimulation (59). State dependence can therefore be assumed to arise from pattern-specific interactions within the recurrent circuit rather than general frequency-dependent adaptation. The difference in performance between networks can be attributed to either varying length in synaptic responses or features in the randomly generated 50 action pairs.

### RL agents increase reward during training

To identify patterns that achieve clockwise firing, we tested different RL agents operating on the continuous and discretised action spaces. While MAB approaches operate state-free, LCBs contain first-order linear state dependency (for a detailed description, see Methods). All agents were tested for 4800 steps (20 min) after being trained for 10k to 120k steps, which is equivalent to 40 min to 8.3 h of interaction with the neuronal network. The agents used PCA with a bin width of 8 samples for state compression (DCNN was tested for a subset but showed no clear performance boost and complicates the fitting, see Fig. S22).

The average reward achieved during the test phase is shown for different networks and different DIVs in Fig. 4a. As discussed above, the reward variance can affect the average reward. Thus, the reward distribution was also analysed by calculating the average across the 20 % highest rewards.

**Fig. 4.**
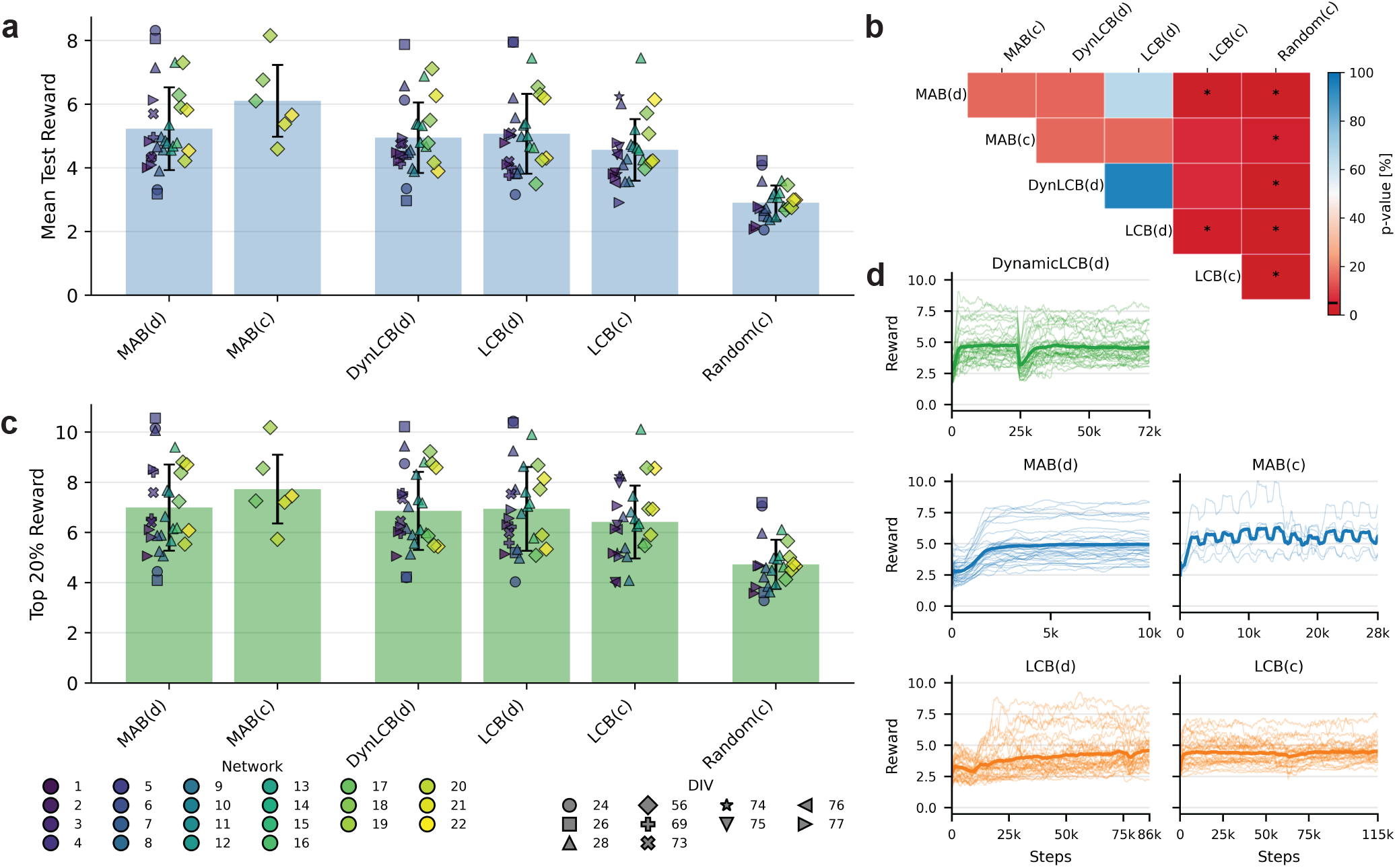
All agents learn to improve reward over random stimulation. **a**, Mean test reward across agents, networks, and DIVs. MAB and LCB are tested on both the continuous (c) and discretised (d) action space. DynamicLCB operates on the discrete space with an additional state-dependent action value matrix. Colour encodes network identity, shape encodes DIV. **b**, Pairwise statistical comparison of mean test reward. Colour encodes the p-value of a two-sided Wilcoxon signed-rank test on min-max normalised, per-network averaged rewards (for details, see agent comparison in Methods). Asterisks denote significance at *α* = 0.05. All agents perform significantly better than random stimulation. Differences between agents are small. The continuous LCB performs worse than the discretised version and the discrete MAB. **c**, Top 20 % test reward, defined as the mean of the highest-scoring 20 % of test samples per run. The corresponding pairwise comparison matrix is shown in Fig. S19. **d**, Training curves for each agent, excluding the random baseline. Thin lines show individual network runs smoothed with a 1,000-step moving average, left-padded with NaN values. Bold lines show the mean across networks. Agents differ in the total number of training steps required for plateauing. For DynamicLCB, the onset of state-dependent action switching is visible at approximately one-third of the training steps. For the adaptive MAB the fluctuations can be seen when the action space is adjusted every 2,500 steps. The continuous LCB shows only a minor increase in average reward over the majority of the training steps. Training with a shorter moving average filtering of 50 steps is provided in Fig. S23 for the first 1,000 steps, the full training, and the transition to the test phase.

A clear difference between networks is visible, as was also shown in Fig. 2 (data separated by network and DIV is shown in Fig. S18). All agents perform significantly better than random stimulation (see comparison matrix in Fig. 4b). Compared to the maximum average reward of the 500 random actions (Fig. 2i), all agents identify in the worst case comparably high-performing policies. The state-based approaches perform similarly well as the state-free.

The continuous MAB that iteratively adapts the action space shows no significantly higher average reward. The LCB agent operating on the discretised action space outperforms the one with the continuous space. Since the discrete action space is a strict subset of the continuous space, this cannot be attributed to the action space itself but must arise from limitations of the learning algorithm. The continuous LCB models reward as a linear function of per-electrode features without cross-terms between the action of each electrode, optimising each electrode’s timing independently. This prevents it from capturing interactions between stimulation timings on different electrodes present in neuronal cultures (16). The discrete agents implicitly handle such relationships by treating each stimulation pattern as an atomic action.

The learning curves in Fig. 4d (see Fig. S23 for more detail) show fast convergence for the MAB, as it is state-free and operates on the discretised action space. During training, oscillations from optimising the precise stimulation timing can be seen for the continuous MAB, with visible drops when opening up the action space. The dynamic LCB variant first trains the action value, effectively mimicking the MAB. When the state is incorporated, performance drops but subsequently recovers when the state-vector *W*_*a*_ is trained. The LCB shows successive improvement across all learning steps. Compared to the dynamic variant, the fitting of the state-vector with poor estimates of the action value leads to lower rewards during learning but similar resulting performance. The continuous version, however, improves performance only in the very beginning.

We also compared the reward associated with the evoked response to spontaneous activity recorded for 5 min before stimulation for one of the MEAs (MEA A). We segmented the spontaneous recording by splitting it when there were no spikes recorded on all four electrodes for 5 ms and discarding chunks with a single spike. The results are shown in Fig. S21. The distribution shows for three of the four networks, higher maximum rewards for stimulated sequences. This could be explained by either additional induced activity or by triggering of multiple spontaneously present sequences, which are then interleaved to maximise reward. The sequences in spontaneous activity could also explain the minimum rewards associated with random stimulation, as the observed rewards are in the same range for both cases.

### Actions achieving maximum reward are non-trivial

After training, the best-performing actions were analysed. The example responses in Fig. 5a confirm response stability even after many hours of continuous stimulation. A video of the best-performing action per agent for an example network is provided in the Supplementary Information (a screenshot is provided in Fig. S24). The histograms in Fig. 5b show that agents exploit the full range of the action space for their best-performing actions. Many electrodes have the highest probability for no stimulation. This could be explained by the fact that with a stimulation, neurons will be activated that then cannot fire again in the recording window. They thus cannot contribute to the reward, leading to lower rewards. The distributions split by network and training run is provided in Fig. S25. No agents systematically collapse onto trivial patterns such as simultaneous stimulation on all electrodes. Different training runs on the same network converge to distinct actions, indicating that multiple local maxima in the reward function exist. The continuous LCB shows a high proportion of minimal delay stimulation, which can be attributed to the per-electrode optimisation, as mentioned above.

**Fig. 5.**
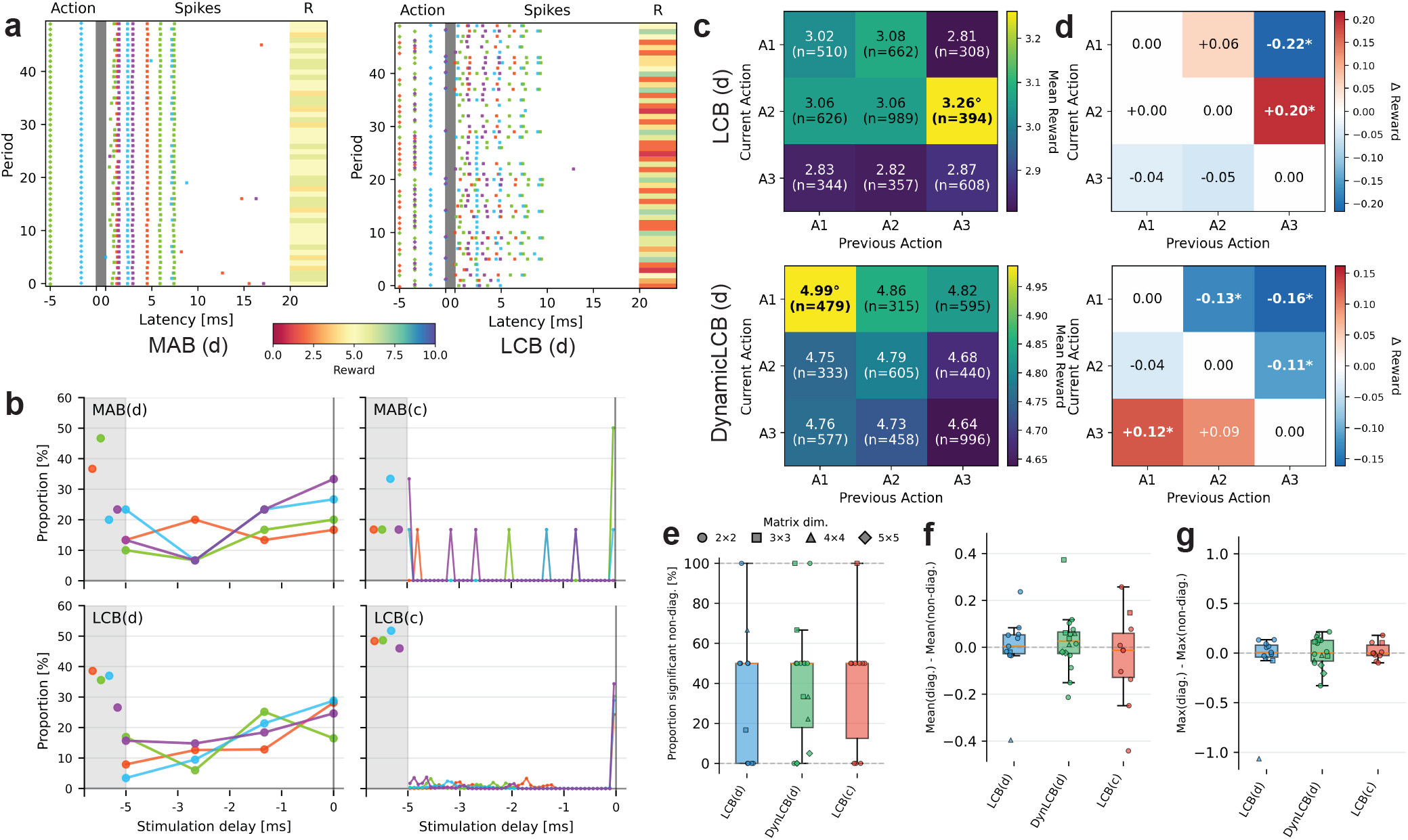
Best-performing actions are non-trivial and state-based agents exploit action switching. **a**, Example post-stimulus raster plots during the test phase on the same network for an MAB (left) and discrete LCB (right) agent, respectively. Stimulation timing is marked with diamonds, spikes are represented by dots, both colour-coded according to Fig. 1a. The reward per period is shown on the right. Responses remain stable even after hours of continuous stimulation. **b**, Distribution of stimulation delays chosen by the respective best-performing action averaged over different networks for four types of agents. A single bin at the left represents no stimulation on that electrode. Actions span the entire action space. **c**, Reward matrix for an agent on one network. Actions were grouped by computing the L2 distance between consecutive four-dimensional action vectors and partitioning these distances into equal-frequency quantile bins (target *n* = 5). Adjacent bins whose width fell below 0.25 were merged. The effective number of action groups varied between agents and networks. Matrix entries were aggregated by current action group (rows) and previous action group (columns). Each cell shows the mean reward and number of trials. The diagonal corresponds to repeated actions, non-diagonal entries to action switches. The agent applied certain action pairs more frequently, indicating learned switching preferences. Bold text and symbols * mark the best-performing actions (average reward significantly higher; Mann-Whitney U, *p <* 0.05). **d**, The matrices show the corresponding reward difference compared to repeating the same action. Asterisks denote statistically significant differences (Mann-Whitney U, *p <* 0.05). Several action switches yielded a measurable reward modulation over repetition. **e**, Proportion of non-diagonal action pairs showing a statistically significant reward difference, summarised across all networks for each state-based agent type. At least some beneficial switching pairs exist for all agents, though the proportion varies across networks. **f**, Difference in mean reward between diagonal (repeated) and non-diagonal (switched) entries. No agent showed a significant bias (two-sided Wilcoxon signed-rank, all *p*_bonf_ *>* 1.0). The effect was centred near zero, indicating that on average the reward benefit from switching was modest, with positive and negative exceptions. **g**, Difference in maximum reward between diagonal and non-diagonal entries. No agent showed a significant difference (two-sided Wilcoxon signed-rank, all *p*_bonf_ *>* 0.05).

Given the goal of inducing clockwise firing, an intuitive expectation would be that optimal actions consist of clockwiseordered stimulation delays. However, the best-performing actions show no preference for clockwise temporal ordering of the stimulation electrodes. This is consistent with the non-trivial relationship between stimulation site and response propagation in microchannel-based networks. Extracellular stimulation activates axons at the electrode site regardless of their projection direction, generating both orthodromic and antidromic propagation (17, 60). Therefore, there exists a multitude of presynaptic and postsynaptic combinations that are not selectively activated. As a result, there can be convoluted probabilistic activation pathways, rendering the mapping from spatiotemporal stimulation pattern to the evoked network response based on the network topology alone challenging. The RL agents must discover effective patterns through interaction.

### State-based approaches can exploit action switching to improve reward

While state-free agents converge to a single best action during exploitation, state-based agents (LCB variants) can alternate between actions conditioned on the previous response. We analysed whether these agents exploit the state dependence characterised in Fig. 3. For each state-based agent, we constructed a reward matrix indexed by the current and previous action group applied during the test phase, where groups were defined by binning the L2 distance between consecutive action vectors into quantile-based bins (Fig. 5c). The diagonal entries represent trials where the same group was repeated, while non-diagonal entries represent switches between groups. The reward difference between these conditions (shown in Fig. 5d) reveals whether switching to a particular action after a specific predecessor yields a higher or lower reward than repeating it. Several state-based agents show statistically significant reward differences, confirming that they learn to exploit action-pair-specific state dependencies. The proportion of significant non-diagonal entries and the associated reward differences across all networks and agents are summarised in Fig. 5eg. While at least some beneficial switching pairs exist for all agents (Fig. 5e), the mean reward difference between repeated and switched actions is centred near zero across agents (Fig. 5f). No agent shows a significant difference in maximum reward between the two conditions (Fig. 5g). This indicates that, on average, the reward benefit from action switching remains modest.

While action switching provides a measurable reward increase for individual action pairs, the overall effect remains modest. As a consequence, the state-based agents do not surpass the performance of the state-free MAB, which benefits from converging to a single high-reward action without the added complexity of learning a state-dependent policy over a limited number of interactions. This is not unexpected with respect to the state dependency analysis in Fig. 3, where not all action pairs showed significant state-dependent responses, despite the experimental design specifically targeting this. The state compression has been further tested for separability of the response. Therefore, it is not known whether it provides the agent with sufficient and relevant cues to inform changing the stimulation. The state definition used here may further be insufficient to describe the network dynamics, which in turn violates the Markov property underlying the MDP formulation. Recordings capture only supra-threshold activity from the microchannels. Further, the 20 ms post-stimulus window captures incomplete information about the ongoing dynamics, considering the timescales of synaptic transmission and the stimulation period of 250 ms. Beyond these limitations, intracellular molecular states, including ion channel distributions, second messenger cascades, and plasticity-related receptor modifications, modulate neuronal responses but remain entirely inaccessible to extracellular measurement.

## Summary and Outlook

Deriving the input-output function of biological neuronal networks remains challenging, even in topologically constrained systems. The space of possible spatiotemporal stimulation patterns grows combinatorially with the number of electrodes and the temporal resolution, with the additional challenge that evoked responses depend on prior stimulation history. Exhaustive open-loop characterisation of this space is intractable, and the non-trivial relationship between stimulation site and response propagation (17, 60) prevents direct mapping of the spatiotemporal stimulation pattern based on the network topology. Addressing this calls for closed-loop experimentation, yet accessible systems that combine deterministic stimulation timing with single-spike-resolution readout have been lacking.

In this work, we addressed this challenge by embedding topologically constrained biological neuronal networks in a closed-loop reinforcement learning framework. RL agents interacted with the networks in real-time, selecting spatiotemporal stimulation patterns based on the observed post-stimulus response and a reward signal encoding a user-defined objective. As a proof of principle, we chose the induction of clockwise-circular firing sequences in engineered recurrent networks. Unlike previous closed-loop approaches that operated on scalar network summaries at timescales of seconds, our system encodes the full spatiotemporal spike pattern into the agent’s state representation and operates at millisecond-scale round-trip times, enabling control at single-spike resolution.

We first established that evoked responses are stable and separable across the action space over hours of continuous stimulation, with approximately 90 % of actions showing stationary reward signals. State dependency was detectable in a substantial subset of action pairs, consistent with intrinsic excitability fluctuations (58) and synaptic dynamics (19). All tested RL agents outperformed random stimulation and converged on non-trivial actions spanning the full action space, rather than mirroring the clockwise target motif. State-based agents learned to exploit the identified state dependence through action switching, yielding measurable reward benefits for specific action pairs. Notably, the platform resolved these state-dependent reward effects at the level of individual action pairs, a sensitivity inaccessible to systems operating on scalar readout functions at the order of seconds. The fact that this advantage did not directly translate into overall performance gains over the state-free multi-armed bandit agents may be explained by not capturing the entire internal state. Furthermore, the operational state representation, as discussed above, incorporates only a temporal subset of the underlying network dynamics. Future work could address this through richer state definitions, for instance by extending the observation window or incorporating multi-step history. It may also be possible to explicitly model the hidden dynamics as a partially observable MDP (POMDP), in which agents maintain a belief over the latent network state rather than treating each observation as complete.

The current technical implementation opens several avenues for further improvement. The electrical stimulation artefact blanks a temporal window around the stimulus, obscuring the directly induced spikes that are among the most reliable components of the evoked response. Combining our framework with optical stimulation methods (61) or high-density MEAs (12) could recover this temporal window and further improve reward signal quality. On a similar note, a substantial proportion of neurons within each compartment are directly activated by the stimulation electrode. Modifications to the microstructure or single-cell seeding techniques (62) could increase the proportion of synaptically driven vs. directly evoked activity, providing richer network-mediated responses for the agent to exploit. Under stimulation with extracellular current pulses, the membrane is either depolarised to threshold, resulting in an action potential, or no detectable response is induced (63). With fewer neurons being directly recruited by stimulation, influences on the subthreshold potential from spontaneous firing and stimulation history would be sustained longer and could therefore increase state dependency (19). Importantly, the modular design of our closedloop system readily accommodates such extensions without changes to the RL framework itself.

The system presented here establishes an accessible, low-cost, open-source research tool for automated, goal-directed functional characterisation of engineered neuronal networks. The approach is not restricted to the clockwise firing objective demonstrated here; any quantifiable network response metric can serve as the reward signal, making the platform a general-purpose tool for mapping input-output functions of biological neuronal networks. Building on our previously published inkube platform (49, 50), the system is entirely based on off-the-shelf components and 3D printed parts, with all manufacturing files provided. With the advanced control of the culture environment, it is possible to run experiments for weeks without human intervention. The platform enables systematic identification of stimulation patterns that evoke specific target responses at single-spike resolution, exploiting efficient closed-loop optimisation. The framework may also be readily applied to biocomputation, where efficient feature encodings and decodings can be applied to derived input-output transformations. Further potential applications include the development of control algorithms for therapeutic electrical stimulation. All design files and code for the closed-loop system and RL agents are publicly available, facilitating adoption and extension by the broader community.

## ACKNOWLEDGEMENTS

This research was supported by ETH Zürich, the Swiss National Science Foundation (SNSF) [project numbers 182779 and 10001282], the Human Frontiers Science Program (HFSP), and the Swiss Data Science Center (SDSC). This work has further been supported by the Center of Living Systems, Chicago and the US National Science Foundation, Division of Physics (PHY-2317138). Stephan J. Ihle is thankful for the support provided by Schmidt Sciences. Furthermore, Stephan J. Ihle is a Chan Zuckerberg Biohub Spoke Awardee.

We thank Giulia Amos for her help with culture preparation and imaging and Joël Küchler for his valuable feedback on the current state of biocomputing approaches, both from the Laboratory of Biosensors and Bioelectronics.

## AI usage statement

Claude (Sonnet 4.5, Sonnet 4.6, Opus 4.5, and Opus 4.6; Anthropic, San Francisco, CA, USA) was used for language editing of the manuscript text and as a coding assistant. All outputs were reviewed, verified, and edited by the authors. No AI tool was used for data collection, interpretation of results, or formulation of scientific conclusions.

## Data and material availability

Design files for manufacturing and a detailed components list are available at https://doi.org/10.3929/ethz-c-000797812 under the Creative Commons Attribution±ShareAlike 4.0 International licence (CC BY-SA 4.0), with a basic components list in Tab. 1. The software for closed-loop stimulation and RL is available on GitHub https://github.com/maurer-lbb/inkube_software.git. Electrophysiology data and analysis scripts are available at https://doi.org/10.3929/ethz-c-000798493.

## CRediT author contributions

**Benedikt Maurer:** Conceptualisation, Methodology, Software, Validation, Formal analysis, Investigation, Data Curation, Writing - Original Draft, Writing - Review & Editing, Visualisation, Supervision. **Vaiva Vasiliauskaitė:** Methodology, Formal analysis, Writing - Review & Editing. **Julian Hengsteler:** Conceptualization, Methodology, Writing - Review & Editing. **Gino Cathomen:** Methodology, Writing - Review & Editing. **Tobias Ruff:** Project administration, Writing - Review & Editing. **Cedric Schmid:** Investigation, Software, Writing - Review & Editing. **János Vörös:** Conceptualisation, Methodology, Writing - Review & Editing, Supervision, Project administration, Funding acquisition. **Stephan J. Ihle:** Conceptualisation, Methodology, Software, Validation, Formal analysis, Investigation, Data Curation, Writing - Original Draft, Writing - Review & Editing, Visualisation, Supervision.

