## Supplementary Information for "A reinforcement learning framework for goal-directed stimulation of engineered neuronal networks"

### Supplementary Note A: Software Architecture

**A.1. System Overview.** The software enables real-time closed-loop control of neuronal networks through a modular architecture that significantly extends the previous implementation (51) (see Fig. S1). Key improvements include: (1) a C++ processing pipeline for low-latency spike detection, (2) ZeroMQ-based inter-process communication for scalable data distribution, (3) USB 2.0 command interface for deterministic stimulation timing, and (4) continuous HDF5 data logging from Python processes. These architectural changes address critical limitations of the previous system. The C++ backend reduces spike detection latency to <2 ms, enabling faster closed-loop responses. The ZeroMQ publish-subscribe pattern allows multiple analysis processes to subscribe to data streams independently without blocking acquisition. USB command transmission provides more reliable timing control compared to UDP-based stimulation commands. Combined, these improvements enable sub-20 ms closed-loop latencies suitable for reinforcement learning experiments.

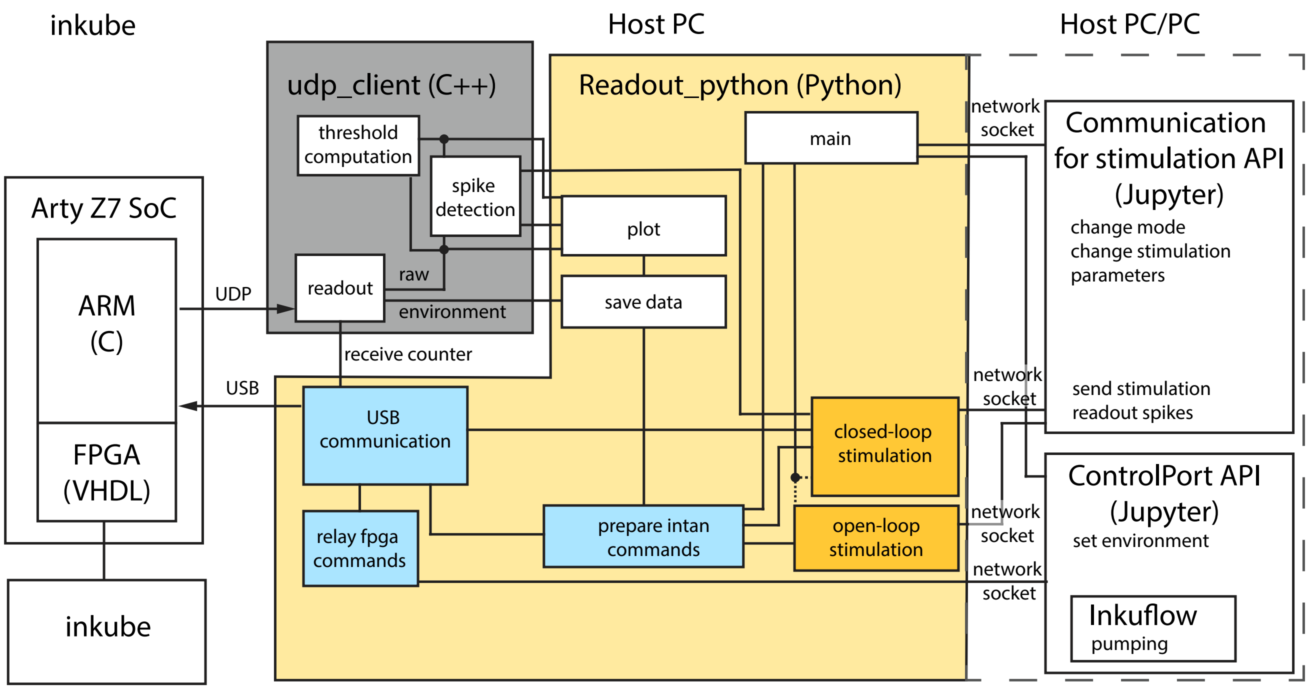


**Fig. S1. Software architecture showing data flow between acquisition, analysis, and control modules.**

**A.2. C++ Processing Pipeline.** The acquisition and spike detection pipeline is implemented in C++ for minimal latency and efficient memory management. The source code is available at (66) (branch: `cpp_backend`). The pipeline consists of four core components that communicate via UNIX domain sockets and ZeroMQ publishers:

**UDP Receiver:** Receives 1428-byte packets from the FPGA at 17.361 kHz containing voltage data from all 240 channels plus environmental sensors. Each packet is immediately distributed to four spike detector processes via UNIX domain sockets (`/tmp/raw_data_socket[0-3]`), with each process handling 60 channels. The receiver runs in a dedicated process to ensure no packets are dropped during high network activity.

**Spike Detectors:** Four independent processes perform parallel spike detection, each processing 60 channels in real-time. Raw voltage data undergoes high-pass filtering (300 Hz cutoff, 2nd-order Butterworth) followed by threshold-based detection. Thresholds are computed as 6× the median absolute deviation of the filtered signal, updated every second with exponential smoothing ($\alpha = 0.25$). When the negative threshold is exceeded, the detector tracks the local minimum over 20 samples (1.15 ms) and extracts a 45-sample waveform (2.6 ms). Channels are then blanked for 25 samples to prevent duplicate detections. Detected spikes are forwarded to the spike collector via internal UNIX sockets.

**Spike Collector:** Aggregates spike events from all four detectors and batches them at 1 ms intervals before publishing via ZeroMQ (port 5556). Each published message contains the spike count followed by packed binary data: channel ID (uint8), package ID (uint32), cycle count (uint16), and waveform (45 float32 values). This batching reduces ZeroMQ overhead while maintaining sub-millisecond temporal resolution for closed-loop control.

**Plot Server:** Consolidates filtered voltage data and detection thresholds from all spike detectors for real-time visualisation. Data is shared (published) via ZeroMQ (port 6001) in 128-sample chunks containing raw traces, filtered signals, and current threshold values for all 240 channels. This enables Python GUI processes to subscribe and render data without impacting the acquisition pipeline.
The C++ implementation achieves deterministic processing latency through: (1) preallocated circular buffers for zero-copy operations, (2) lock-free inter-process communication via UNIX sockets, (3) SIMD-optimised filtering operations, and (4) isolated processes to prevent scheduler interference. Total spike detection latency from UDP packet arrival to ZeroMQ publication is consistently <2 ms on the host system, a Lenovo ThinkStation Tiny P360 running on Linux Ubuntu 22.04 LTS.

**A.3. Inter-process Communication.** Data sharing between modules uses ZMQ sockets to enable low-latency communication while maintaining modularity. This architecture allows independent development and real-time performance optimisation of individual components.
The system employs a hybrid socket architecture optimised for different communication requirements (Fig. S1). Internal communication between C++ processes (UDP receiver, spike detectors, plot server, spike collector) uses UNIX domain sockets for minimal latency.
External data distribution to Python analysis processes uses ZeroMQ TCP sockets. Separate ZMQ publishers provide real-time access to: (1) detected spike events with waveforms (port 5556), (2) raw package data (port 5557), (3) filtered voltage data of all channels (port 6001), and (4) environmental sensor data, including temperature (port 5551). This architecture enables independent subscribers to access specific data streams without blocking the acquisition pipeline.
The modular design allows spike detection to run in parallel across four independent processes, each handling 60 channels of the 240 total channels. Spike events accumulate in batches with configurable intervals (default 1 ms) before transmission to minimise communication overhead while maintaining real-time responsiveness for closed-loop control.

**A.4. USB Communication Protocol.** The system employs a bidirectional communication architecture that separates high-bandwidth data acquisition from low-latency command control. While voltage data streams from FPGA to PC via Ethernet (UDP) as in the previous version (51), stimulation commands and configuration registers are transmitted in the reverse direction via USB 2.0. This asymmetric design optimises both throughput and real-time responsiveness: UDP handles continuous 240-channel data acquisition at 17.361 kHz without acknowledgement overhead, while USB provides reliable command delivery with hardware-level timing guarantees.
Control commands are transmitted via USB 2.0 bulk transfers using a custom protocol with Vendor ID `0x33FF` and Product ID `0x1234`. Communication is handled through the `libusb` library via Python's `usb.core` interface, operating on endpoint `0x01` for outbound transmission.
Each USB packet transmits up to 64 stimulation commands in a fixed 264-byte structure. Commands begin with an 8-byte preamble (`0x01020304fdfeff00`) followed by the payload specifying stimulation timing, electrode selection, and amplitude parameters. The command structure supports sub-millisecond timing resolution by encoding package IDs that reference the FPGA's internal timestamp counter. During system initialisation, the host first communicates the UDP port assignment to the FPGA through this USB interface, establishing the complete bidirectional communication link. Commands are buffered in Python and transmitted in batches to minimise USB transaction overhead while maintaining <10 ms command latency for closed-loop control.

**A.5. Data Structure and Storage Format.** Experimental data is stored in HDF5 format (Hierarchical Data Format version 5), providing efficient storage and random access for large-scale electrophysiology recordings. The file structure organises data into hierarchical groups with appropriate chunking and compression for different data types.
**Spike Data:** Detected spikes are stored in a dataset `/spikes/events` with compound datatype containing: channel ID (uint8), package ID (uint32), cycle count (uint16), and 45-sample waveform (float32 array). The cycle count is used to account for the reset of the package count every 90 s.
**Raw Voltage Data:** Continuous voltage recordings are stored in `/raw_data/voltage` as a 2D dataset with dimensions [n_samples, n_channels]. Each row represents one package ID containing voltage measurements from all 240 channels as 16-bit signed integers. The dataset uses chunk sizes of (1000, 240) samples to optimise sequential access patterns typical in neural data analysis. Package IDs are stored separately in `/raw_data/package_ids` as a 1D uint32 array for temporal reconstruction.
**Metadata:** Recording parameters are stored as attributes on the root group, including: sampling frequency (17.361 kHz), number of channels (240), electrode layout, stimulation parameters, and experimental conditions. Environmental data (temperature, humidity) is stored in `/environment/temperature` with timestamps in `/environment/timestamps`.
The Python-based H5Saver class (`h5_saver.py`) subscribes to ZMQ endpoints and continuously writes data during acquisition. Separate threads handle spike data and raw voltage data to prevent buffer overflow, with configurable write intervals to balance disk I/O and memory usage. File sizes typically range from 1-5 GB per hour of recording, depending on network activity.

**A.6. Example usage of the RL Gym.** An example of the training of the state representation is shown in the code sample below:

```
env = RealNetworkContinuous(
   action_dim=action_dim,
   stim_length=int(5e-3*fs),
   state_dim=state_dim,
   circuit_id=circuit_id,
   reward_object=ISIseqReward(),
   state_object=DynamicStatePCA()
)
state, _ = env.reset()

spikes = []
elecs = []

for i in range(num_samples):
   action = env.action_space.sample()
   state, reward, terminated, truncated, info = env.step(action)

   spikes.append(info['spikes'])
   elecs.append(info['elecs'])

env.state_object.fit(spikes, elecs)
```

### Supplementary Note B: Hardware System

The hardware system integrates temperature control, electrophysiology recording, and environmental regulation in a compact MEA-compatible platform. The modular design allows independent optimisation of thermal, electrical, and fluidic subsystems.

**B.1. MEA Holder and Temperature Control.** The MEA holder integrates a temperature-controlled base PCB with a heat-conductive region where the glass MEA sits, secured by an MEA frame board (see Fig. S2). A PT1000 sensor provides temperature feedback to a PI controller implemented on the SoC. Bidirectional temperature adjustment uses a thermoelectric element with a heat sink mounted below the control region. The holder connects to the electrophysiology board ground via a soldered cable connection.

The base PCB mounts on a 3D-printed PLA holder with M4-mounted clips on each side for securing the electrophysiology board.

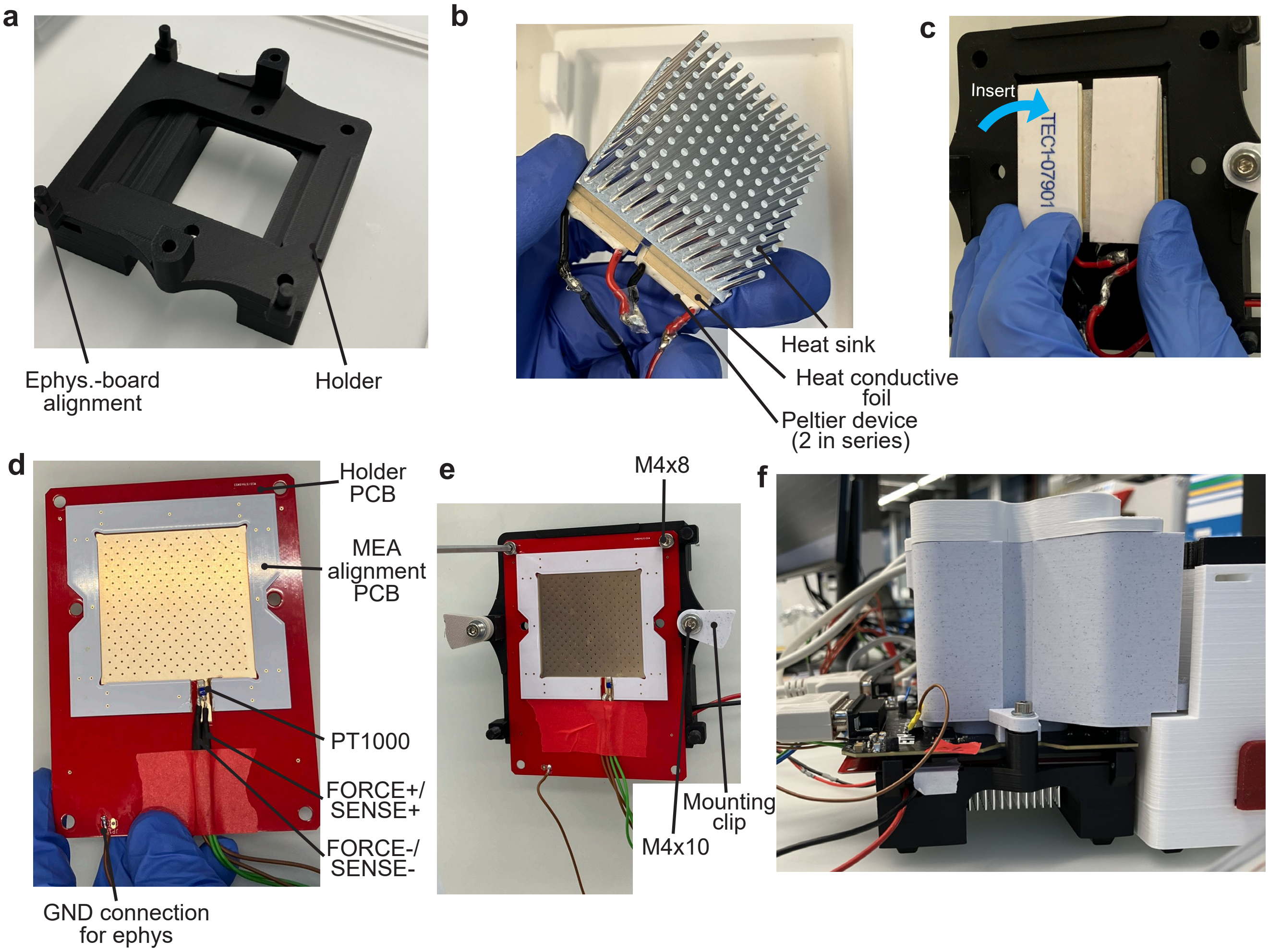


**Fig. S2. MEA Holder assembly.** **a**, The MEA holder is 3D printed with PLA. All holders are equipped with M4 thread inserts. **b**, The temperature actuator is assembled with two Peltier devices in series, and a heat sink, connected with heat conductive foil that is adhesive on both sides. **c**, The temperature actuator is placed inside the holder. The two wires are guided out through the hole on the side. **d**, The holder PCB is manufactured at 0.8 mm thickness, ENIG finish, and with a heat conductive region where the MEA is placed. It is prepared by placing and glueing a PT1000 temperature sensor on the heat conductive area. For a 4-wire impedance measurement, two connections are soldered to each of the leads, one to force the current and one for voltage sense. The wires can be secured with a piece of tape. A gap is cut into the MEA alignment PCB to host the PT1000 before assembly. A wire is soldered to the GND connection for grounding with the electrophysiology board. **e**, The holder PCB is then screwed onto the holder for good thermal coupling. The two mounting clips are also mounted. **f**, Side view after assembly of the chamber.

**B.2. Electrophysiology Board.** Custom electrophysiology boards are shown in Fig. S3. They provide 60-channel recording and stimulation capability using Intan RHS2116 ASICs (Intan Technologies, Los Angeles, CA, USA). The reference electrode connects to ground, with supply voltage distributed through an internal PCB layer. Pogo pin mounting follows the protocol described in Maurer et al. (51).

**B.3. Environmental Chamber.** A 3D-printed PLA chamber mounts on the MEA and connects the medium reservoir to the recording environment. An integrated fan maintains temperature equilibration across the chamber volume. To prevent electrical

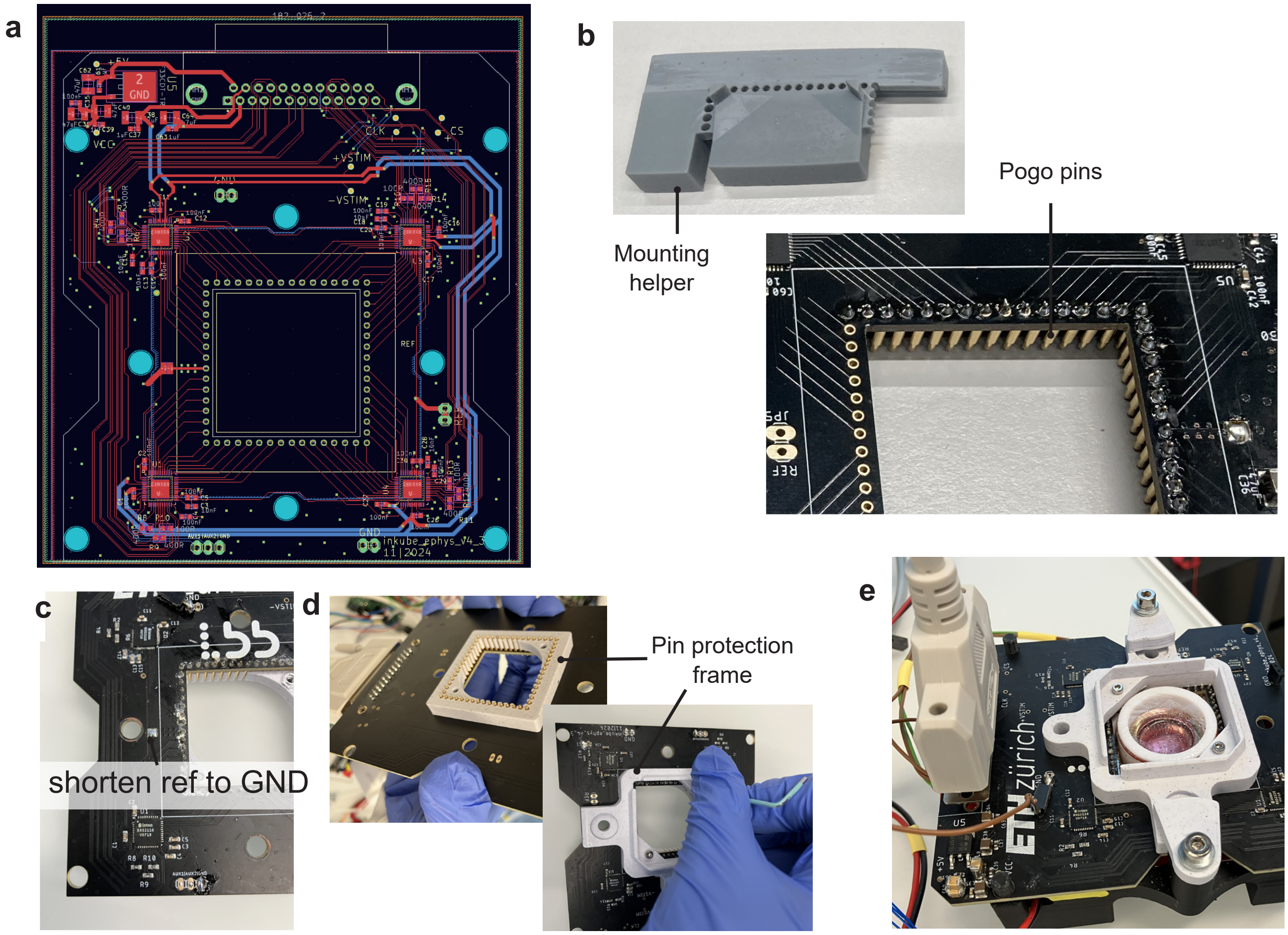


**Fig. S3.** **a**, Electrophysiology board layout. A 4-layer board is required with top and bottom signal layers, one GND, and one $V_{CC}$ layer. **b**, Spring contacts are mounted on the PCB using the mounting helper. **c**, The reference electrode is connected to GND by shorting the two exposed pads. **d**, The pin protection frame is mounted with two M2x12 screws. **e**, The signal DSUB-25 cable connects the electrophysiology board to the SoC shield. The mounting board GND is connected to a GND pin.

noise induction from the fan motor, a custom laser-cut copper housing can be added that provides electromagnetic shielding.

**B.4. Bill of Materials.** The recording system extends the inkudock incubator and inkube electronics presented in (53) and (51), respectively, with a glass MEA-compatible holder, electrophysiology board, and environmental chamber. Table 1 lists the components introduced in this work. For the SoC, SoC shield, power board, and inkudock, refer to (53). PCB design files, 3D CAD models, and detailed component-level BOMs are available in the ETH Research Collection.

| ID | Name | Specification | Manufacturer / Supplier |
|---|---|---|---|
| *Printed Circuit Boards* | | | |
| PCB1 | Electrophysiology board | 4-layer, inkube_ephys v4.3 | Custom, see Research Collection |
| PCB2 | MEA base holder | 2-layer, inkube_base_holder v4.1 | Custom, see Research Collection |
| PCB3 | MEA frame | 2-layer, inkube_MEAholder v4.1 | Custom, see Research Collection |
| *3D-Printed Parts (PLA)* | | | |
| CAD1 | Bottom pin frame | PLA, 3D printed | In-House |
| CAD2 | Top counter frame | PLA, 3D printed | In-House |
| CAD3 | Glass MEA chamber | PLA, 3D printed | In-House |
| CAD4 | Mounting helper | PLA, 3D printed | In-House |
| *Key Electronic Components* | | | |
| IC1 | Intan RHS2116 | Stim/recording ASIC, QFN-44 ($\times$4) | Intan Technologies, USA |
| IC2 | LF33CDT-TR | 3.3 V LDO, TO-252 | STMicroelectronics |
| J1 | D-Sub connector | 182-025-2, 25-pos, RA, female | NorComp / Mouser |
| SC | Spring contacts | 7913-0-15-20-77-14-11-0 ($\times$60) | Mill-Max / Mouser |
| *Thermal Management (per holder)* | | | |
| TD | Peltier device | TEC1-07901, 50$\times$20 mm ($\times$2, series) | Wellen Technology Co., China |
| HS | Heat sink | ICK S 45$\times$45$\times$20, pin type | Fischer Elektronik, DE |
| HF | Thermal foil | WLFT 404, 40$\times$40 mm | Fischer Elektronik, DE |
| T | PT1000 sensor | NB-PTCO-182, 2.0$\times$4.0 mm | TE Connectivity / DigiKey |
| F | Fan | RND 460-00087, 30$\times$30$\times$10 mm, 5 V | Distrelec AG, CH |
| *Consumable* | | | |
| MEA | Glass MEA | 60MEA500/30iR-Ti-gr | MCS GmbH, DE |

**Table 1.** Bill of materials for the glass MEA recording unit introduced in this work. For the SoC (Arty Z7-20), SoC shield, power board, and inkudock incubator, see (53).

**B.5. MEA mounting.** The mounting of an MEA is shown in Fig. S4. To record signals from the MEA, it has to be mounted on the holder, with the reference electrode aligned to the left of the electrophysiology board. The board is aligned to the holder, pushed down, and clamped onto the MEA holder by rotating the clips. Then, the chamber can be mounted by sliding it down into place and optionally tightening the top screw. Finally, the chamber lid is placed.

**B.6. Temperature Calibration.** Medium temperature control requires calibration between the MEA holder sensor and actual medium temperature. Calibration measurements with a reference sensor placed directly in the medium under the water compartment cap established that a holder sensor reading of 32.3 °C corresponds to ≈35.5 °C medium temperature. This offset of 3.2 K accounts for thermal gradients through the MEA substrate and medium layer. The low temperature is ensured to avoid overheating the cell culture in case the water layer fully evaporates.

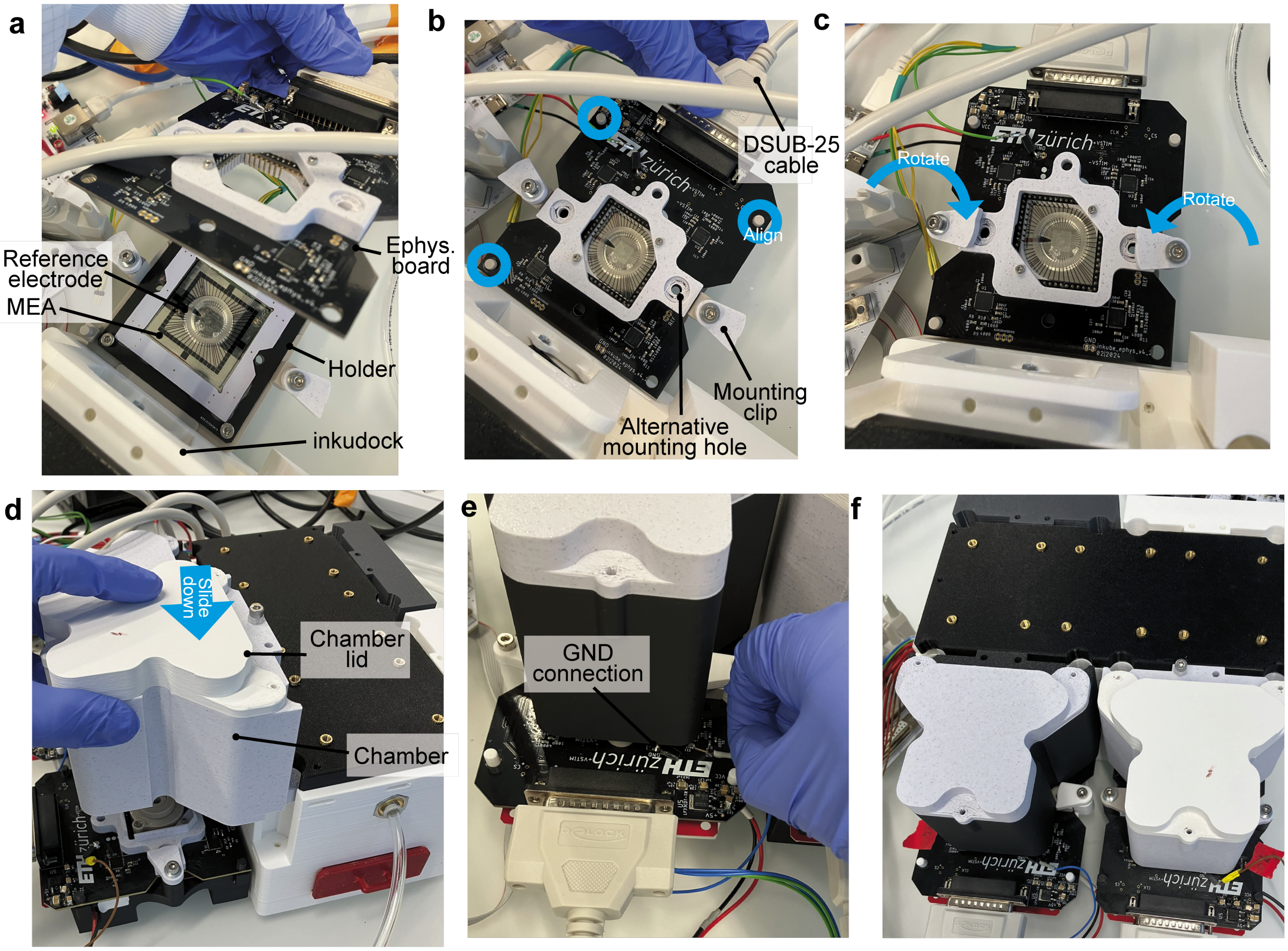


**Fig. S4. MEA mounting and chamber assembly.** **a**, The MEA is placed in the holder board. The reference electrode must be aligned to the left. **b**, The electrophysiology board is connected via the DSUB cable and placed on top of the MEA, aligned by the pins of the holder going into the board. **c**, The board can then be gently pushed down against the force of the pogo pins and the mounting clips are turned to clamp the board. **d**, To seal the MEA off from the lab environment and merge it to inkudock, the chamber is used as a connector. The chamber is slid down until it aligns with the inner ridge of the mounting frame. **e**, The electrophysiology board must be connected with the GND of the holder PCB. **f**, The top view shows two fully mounted chambers with electrophysiology boards.

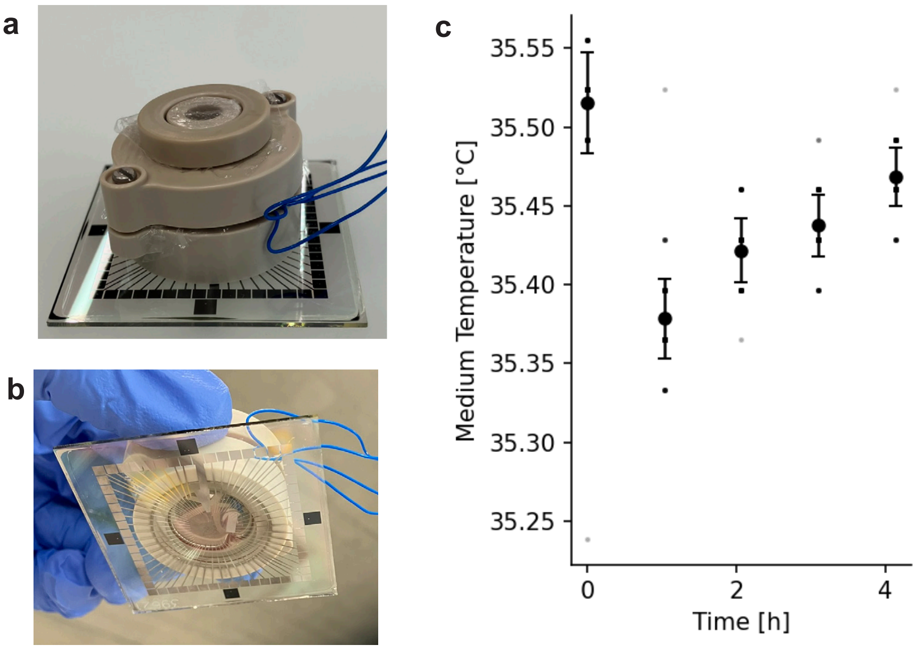


**Fig. S5. Temperature calibration for MEA holder.** **a**, A PT1000 temperature sensor is introduced below the water compartment. A gap is left for the cable. **b**, The sensor measures the temperature from the medium. **c**, Measurements after approximately 24 h of the cap inside the reservoir environment and 5 h after refilling the water compartment and readjusting the MEA holder temperature. The deviations could be caused by ongoing evaporation from the water compartment, as it cannot be sealed airtightly with the sensor in place.

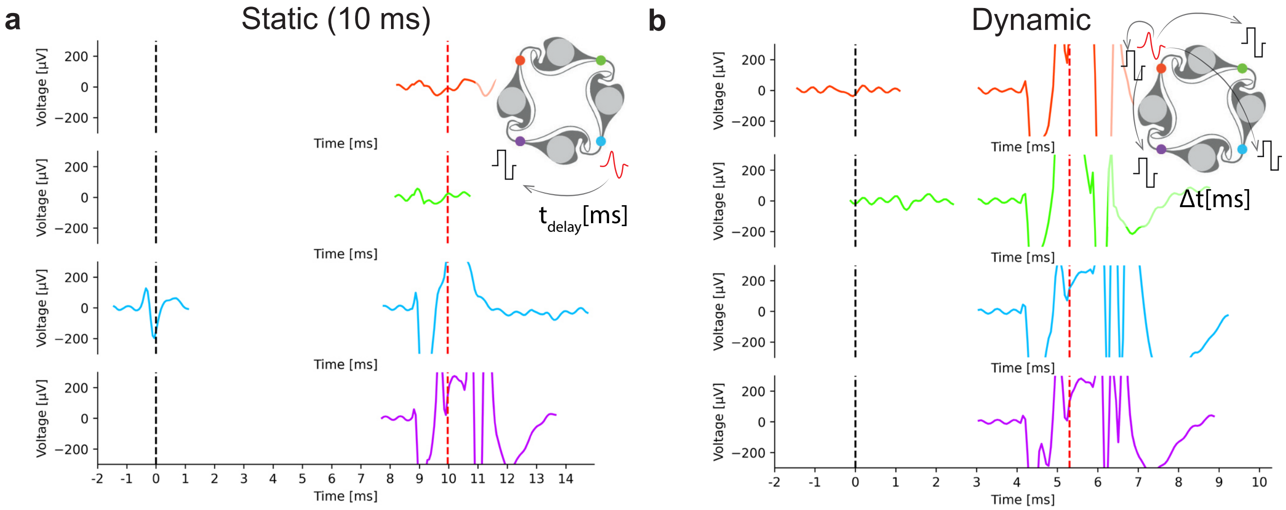


**Fig. S6. Spontaneous spike-triggered stimulation. a** Static mode, where a stimulus with execution timepoint exactly $t_{\text{delay}} = 10$ ms after a detected spike is triggered. Here, a stimulus on the purple electrode is triggered by a spike on the cyan electrode. **b** Dynamic mode, where the stimulus is executed as fast as possible, reducing the time between the triggering spike and the stimulation to about $\Delta t \approx 5$ ms. The relative stimulus alignment $\Delta t$ is consistent between electrodes selected for stimulation. Here, all electrodes are stimulated after a spike on the red electrode.

### Supplementary Note C: Experimental Data

**C.1. Closed-loop minimal round-trip time.** To demonstrate the minimal round-trip time, we trigger a stimulation after the detection of a spike. The schematic of which electrodes are triggered and the time series with the detected spike wave cut-outs are shown in Fig. S6. The round-trip time, including spike detection, internal process communication, and USB transmission, sums to about 5 ms (dynamic mode). When precise timing is desired (static mode), a buffer should be added. Here we show 10 ms total time to account for variation in the transmission time of the USB commands.

**C.2. Stimulation amplitude sweep.** Two amplitude sweeps were performed to cover the full range from 10 to 1,500 nA. In the first sweep, the amplitude was increased from 10 to 500 nA and decreased again; in the second, amplitudes of 50, 500, 1,000, and 1,500 nA were applied in descending order. At each amplitude, a sequence of 200 stimuli that was generated at the beginning of the sweep was applied 20 times at 4 Hz, yielding 4,000 stimuli per amplitude. Post-stimulation time histograms were extracted per electrode and averaged across episodes and stimuli. For amplitudes covered by both sweeps (50 and 500 nA), histograms were averaged across the two measurements before display.

**C.3. Stimulation histogram analysis.** To characterise the temporal precision of stimulus-evoked responses, spike trains recorded across all occurrences of each stimulation pattern were aggregated into per-electrode probability histograms at single-sample resolution (Fig. S8). Response peaks were identified using a local-maximum detection algorithm on the raw probability traces, retaining only peaks whose 0.5 ms moving-sum probability exceeded 10%. Patterns exhibiting no detected peak beyond 10 ms post-stimulus were excluded from further analysis. The relationship between peak latency and peak probability was assessed via Pearson correlation across all retained peaks (Fig. S9).

**C.4. Spike count to reward variability correlation.** Per-action average spike count versus reward standard deviation for all 500 actions across all networks (data from Fig. 2g-h) is shown in Fig. S10. Each point represents one action, colour-coded by network identity (same colour map as Fig. 2i-j). A Pearson correlation analysis revealed a positive correlation between spike count and reward variability, indicating that actions eliciting more spikes tended to produce more variable rewards.

**C.5. Control MEA without cells.** To test the influence of the artefact on the reward metrics, an MEA without cells was tested. As the microstructure is known to have an effect on the artefact spread, it was mounted on the MEA. Stimulation parameters were kept constant. Circuits, meaning the circular structure that would usually form the network, with fully covered electrodes were discarded. A few raster plots generated as in Fig. 2c are shown in Fig. S11. The events occur at around 0.5 and 2 ms, separated by the blind duration after spike detection. To counteract this, the spike detection could be changed to a waveform-based convolutional kernel, or the IIR filter could be adjusted to mitigate ringing.
The quantitative post-stimulation histogram of 500 randomly generated actions as in Fig. 2a is shown in Fig. S12 for four different circuits without cells.
To analyse how much the stimulation artefact can influence the agents searching for the best action, we also computed the reward for the 500 actions. The distribution and standard deviation are shown in Fig. S13. The maximum rewards go as high

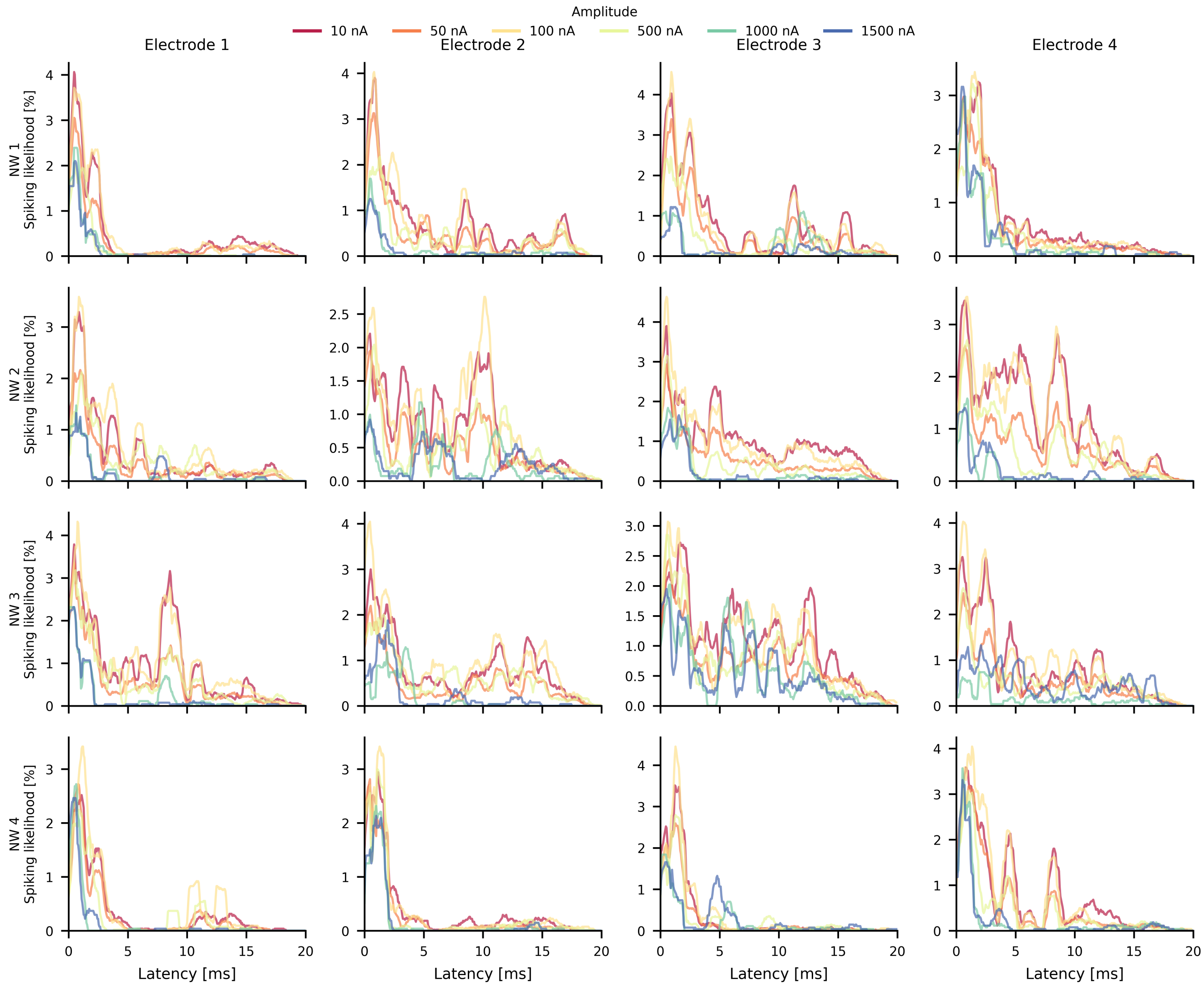


**Fig. S7. Spiking likelihood after a stimulus with different amplitudes.** Post-stimulation time histograms are shown for each electrode and network across the full amplitude range from 10 to 1,500 nA. In principle, a low stimulation amplitude is desirable in order to minimise charge injection. Already at 50 nA, a sufficient proportion of late responses is visible. This low required amplitude can be attributed to the confining effect of the microchannels (67).

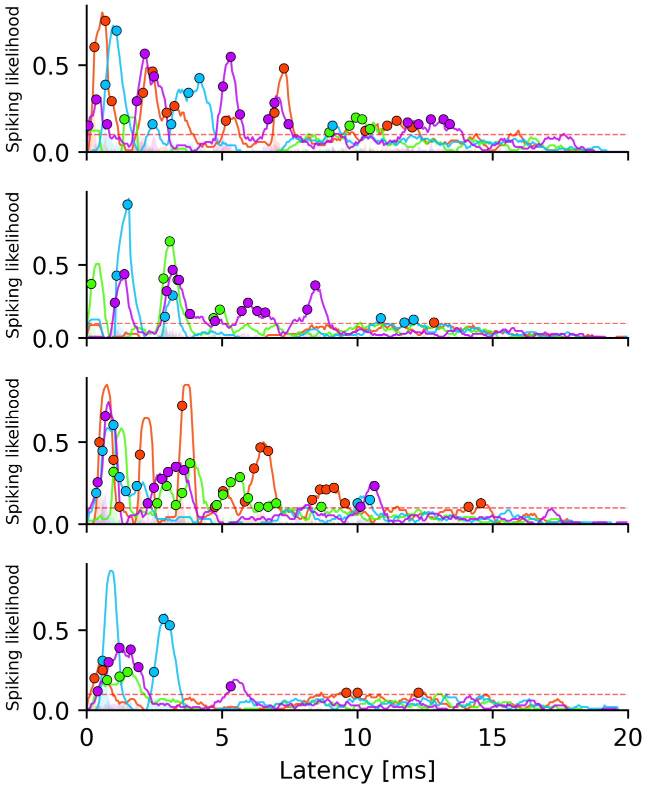


**Fig. S8. Example spike likelihood traces used for jitter analysis.** Post-stimulus spike likelihood for four randomly selected actions. Shaded areas show the raw per-sample spike probability; solid lines show the 0.5 ms moving sum used for peak detection. Colours indicate recording electrodes. Dots mark detected peaks (moving-sum threshold: dashed red line). Peak latencies were extracted from these traces across all actions and networks.

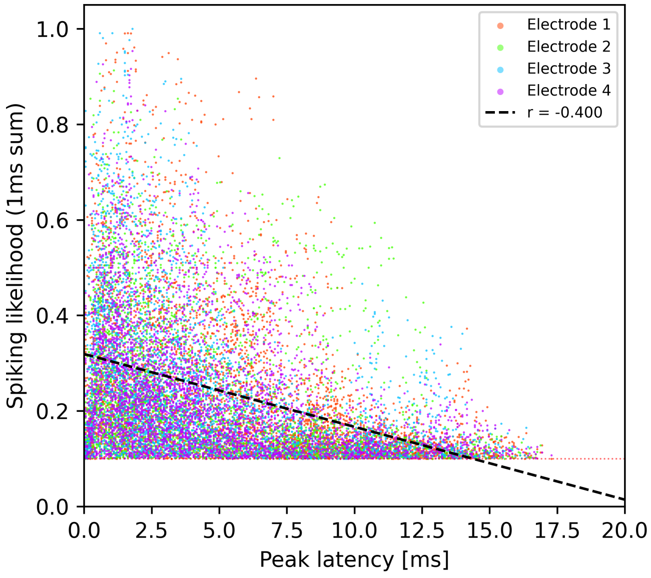


**Fig. S9. Stimulus-evoked response probability as a function of peak latency.** Each point represents a single detected response peak from one stimulation electrode (colour-coded) and one stimulation pattern ($n = 383$ patterns, representative network). Spiking likelihood is quantified as the 0.5 ms moving-sum of the per-electrode spike probability histogram. The dashed line indicates a linear regression fit across all electrodes (Pearson $r = -0.40$), demonstrating that high-probability responses are confined to short latencies. Only patterns containing at least one peak beyond 10 ms post-stimulus were included. The dotted red line denotes the 10 % detection threshold.

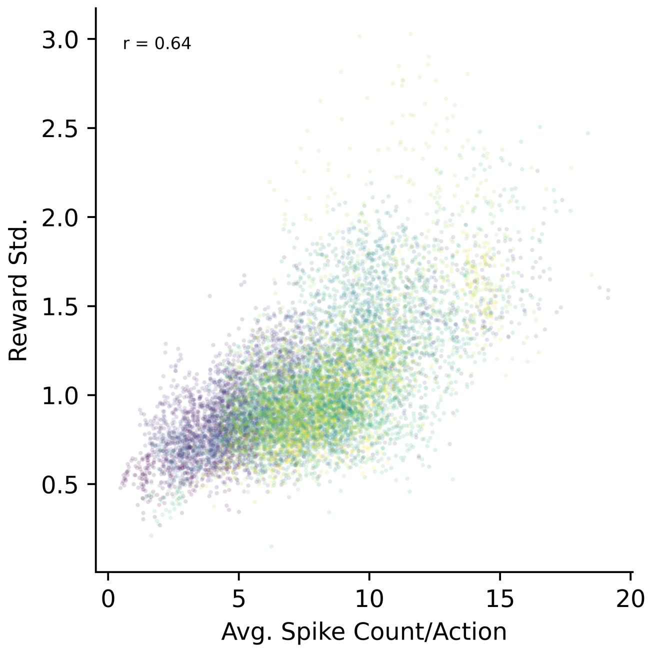


**Fig. S10. Relationship between spike count and reward variability.** Per-action average spike count versus reward standard deviation across all networks. Each point represents one of the 500 actions per network, coloured by network (21 networks). Pearson $r = 0.64$, $p < 0.001$ (two-tailed t-test).

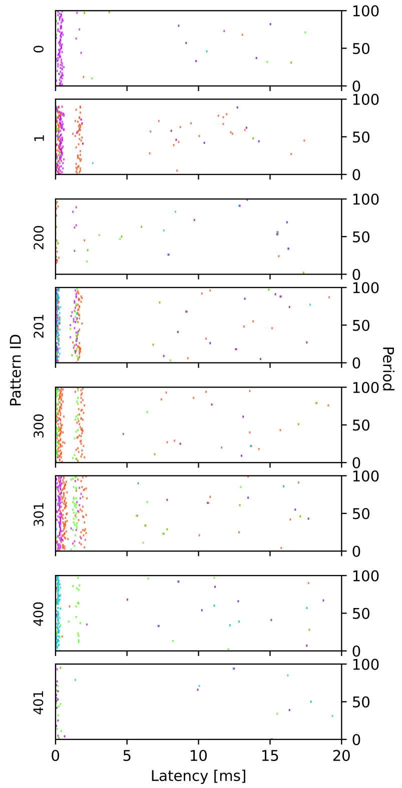


**Fig. S11. Raster plot for MEA without cells showing stimulation artefact.** Raster plot generated as in Fig. 2c for MEA without cells. The raster shows detected events exceeding the spike detection threshold induced by stimulation artefacts.

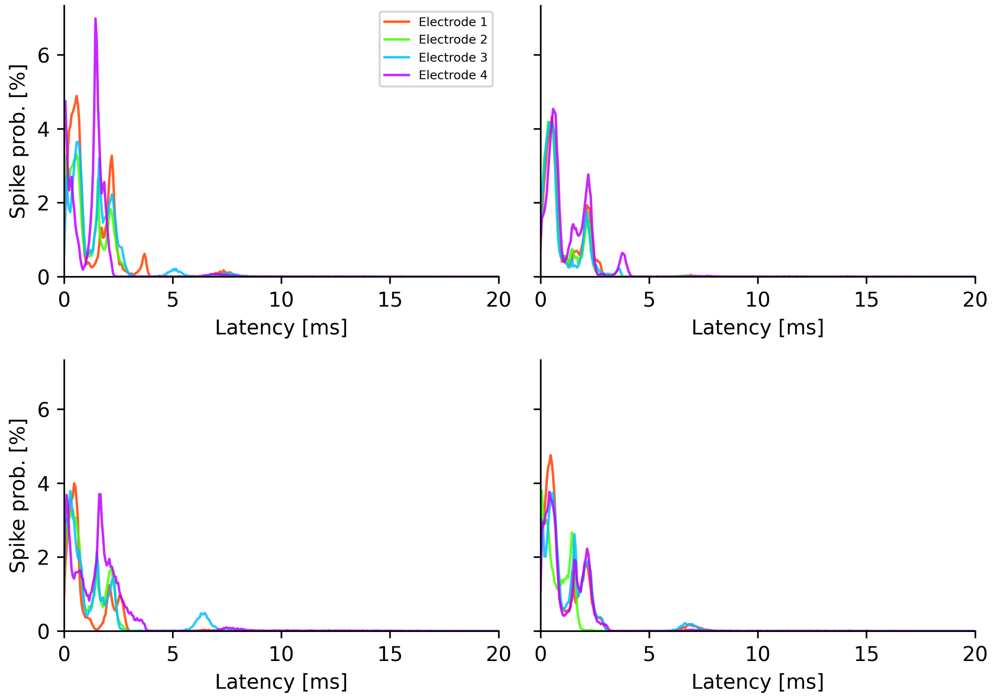


**Fig. S12.** Spike likelihood for the 500 random actions without cells for four different networks. Only some actions induce an artefact. Some artefacts

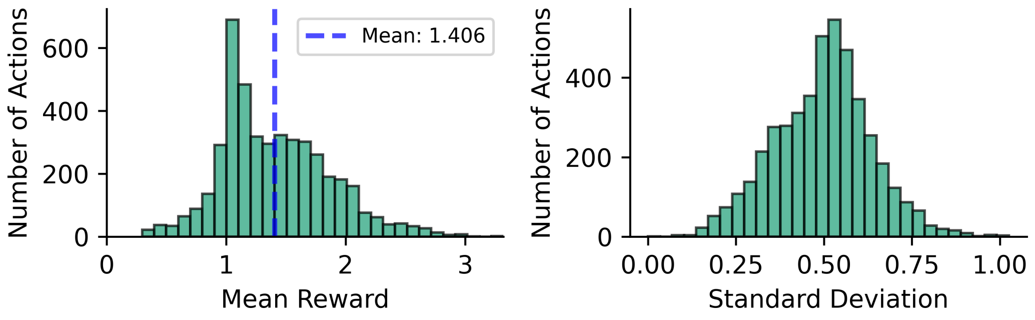


**Fig. S13. Reward statistics for stimulation artefacts on MEA without cells.** The distribution has a large peak at 1, which is the result of no sustained artefacts. Some artefacts lead to rewards as high as 3. These could be exploited by the agents. The standard deviation is much smaller than with the cell culture, showing that there are probabilistic synaptic connections and axonal recruitment at play in the biological networks.

as 3, which is caused by the low minimum time window threshold of the reward metric. The reinforcement agents are able to exploit those, especially because of the low variability. To counteract this, the reward metric could be altered or the blanking time after stimulation increased.

**C.6. State dependency control.** The state separation for distributions with the same current but different previous actions was also tested with a lag of 5 and 10. The bar plot in Fig. S14shows the data from lag 1 that is also presented in Fig. 3e next to the control lags 5 and 10.

**C.7. Agent performance comparison.** Fig. S15shows the mean test reward for each agent broken down by network and recording day, complementing the aggregated comparison in Fig. 4a. Performance varies across networks but the relative ranking of agents is largely consistent. Fig. S16reports pairwise statistical comparisons for the top 20 % of rewards, mirroring the analysis in Fig. 4b. Fig. S17shows the action discriminability index per network, defined as the interquartile range of the 500 per-action mean rewards divided by the median per-action standard deviation. Values above one indicate that differences between actions exceed typical trial-to-trial variability, confirming that the reward landscape is sufficiently structured for agent learning.

**C.8. Spontaneous vs stimulated sequences.** To get a reference point for the limit of the length of stimulated sequences, we extracted the reward during spontaneous recording. We segmented it by splitting when no electrode recorded activity for 5 ms.

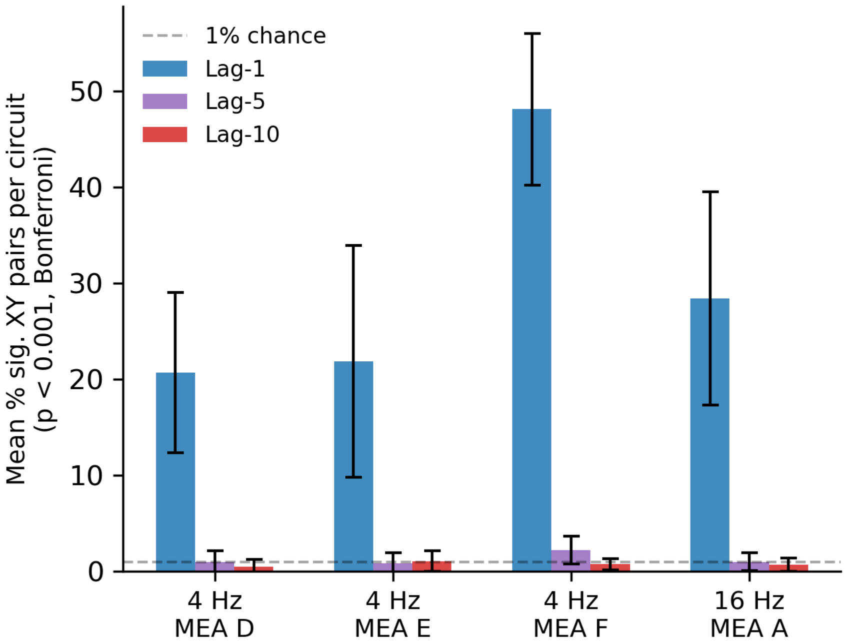


**Fig. S14.** State dependence on the previous action with lag 1 (blue), lag 5 (purple), and lag 10 (red) for PCA-5 (bin = 17 ms) across four recordings (MEA D, E, F at 4 Hz; MEA E at 16 Hz). Error bars show standard deviation across circuits. The dashed line indicates the 1 % significance rate expected by chance. At lag 5 and 10, no individual circuit showed a significant elevation above the 1 % chance level (one-sided binomial test per circuit, FDR-corrected across all circuits and recordings, all $p > 0.01$).

Chunks with at least 2 spikes were considered. The spontaneous activity was recorded for 5 min before the first stimulation. The distributions show much shorter sequences for spontaneous activity, which suggests that the high reward patterns during stimulation contain a combination of activated spontaneous pathways.

**C.9. State compression comparison for agent performance.** Both the DCNN and PCA state compression with a bin size of 8 samples were used with a state dimension of 5 for the agents. The results are shown in Fig. S19. Despite differences in separability of responses in Fig. 3e, there was no observable performance boost for one of the methods with respect to the other.

**C.10. Training data for all agents.** The data from Fig. 4d is shown in Fig. S20 with a shorter window for moving average filtering.

**C.11. Best performing action video.** Fig. S21 shows a screenshot of the video showing the best-performing test action for each agent.

**C.12. Test action distribution.** The data provided in Fig. 5b is shown in Fig. S22 separated by DIV and network.

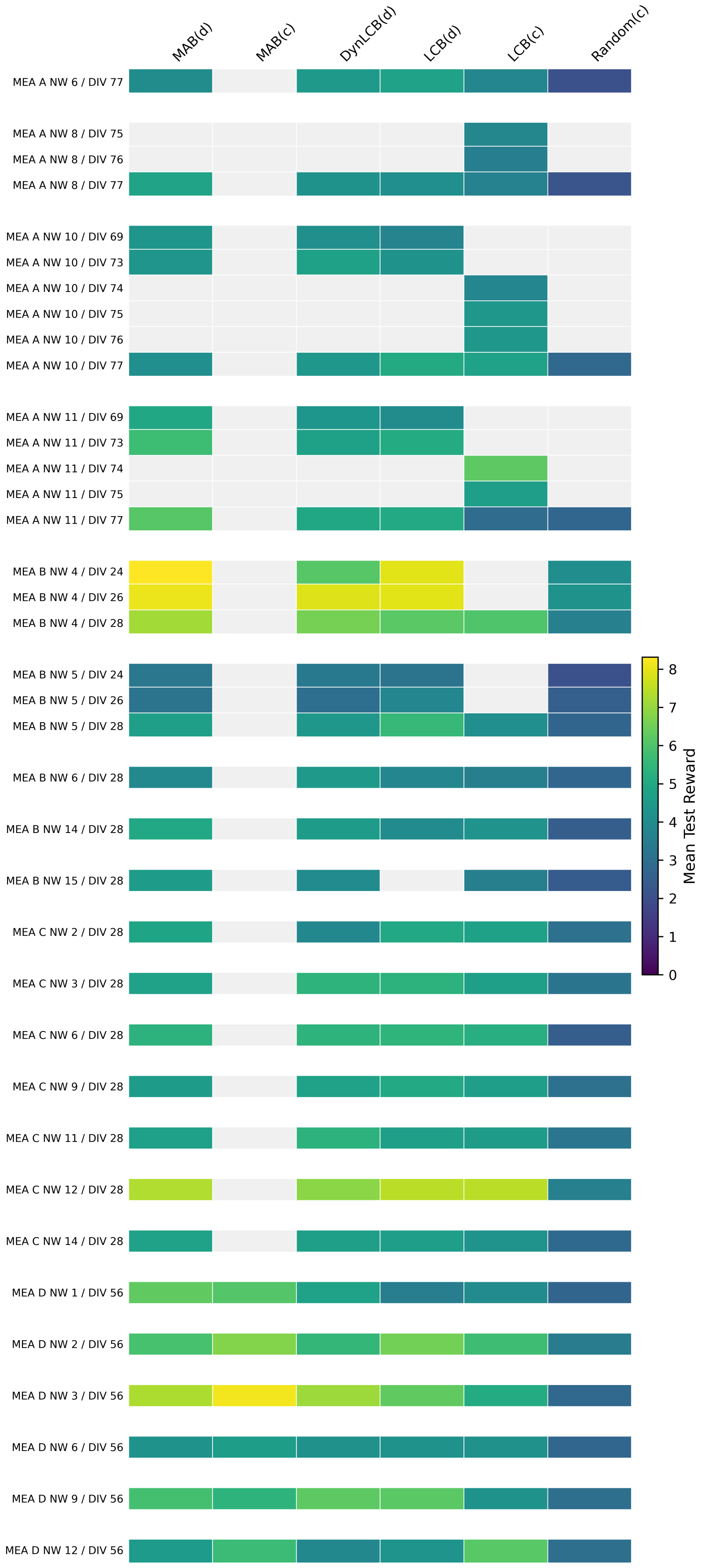


**Fig. S15. Heat map for the data from Fig. 3a separated by network and day.**

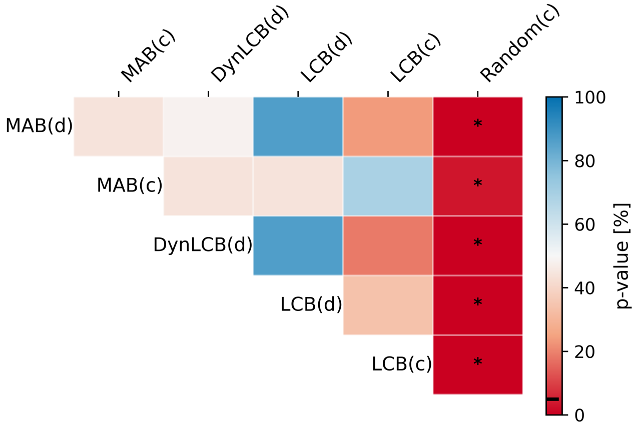


**Fig. S16. Pairwise comparison as in Fig. 3b for the data in Fig. 3c.**

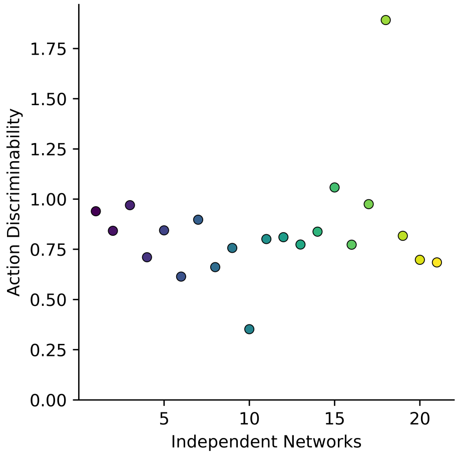


**Fig. S17. Action discriminability index per network.** IQR of per-action mean rewards divided by the median per-action standard deviation. Each point represents one network.

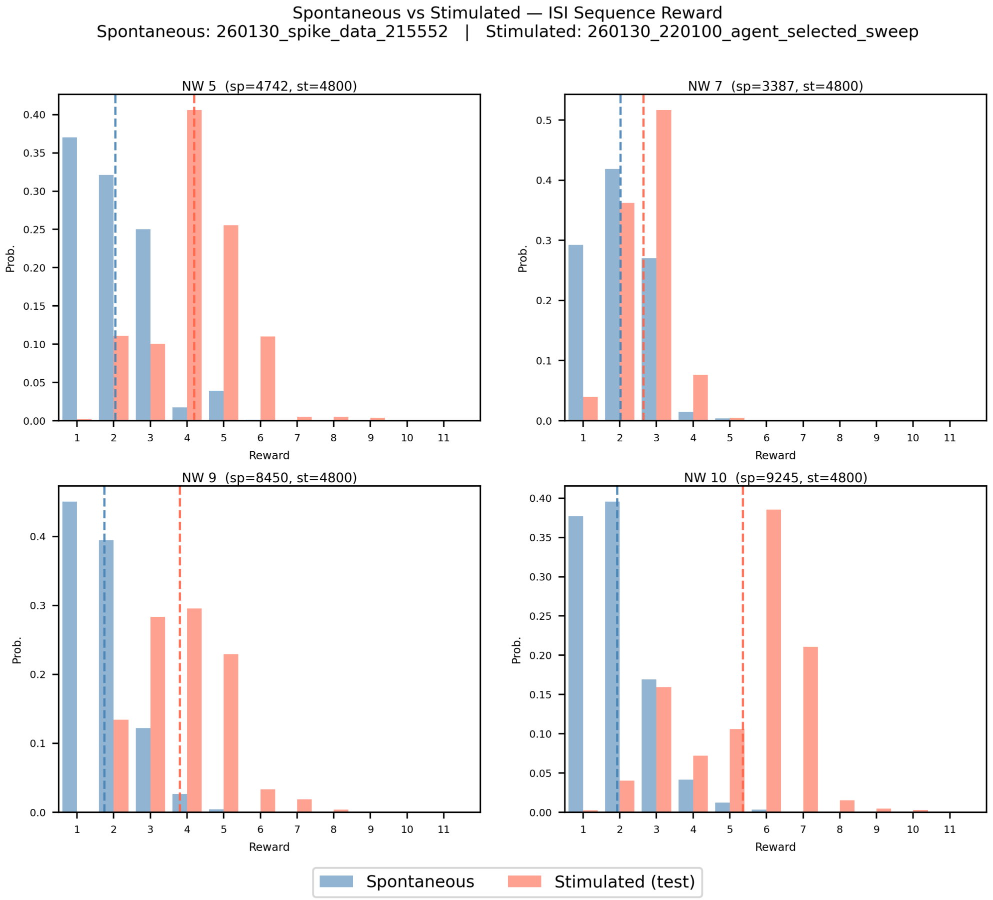


**Fig. S18. Reward comparison of stimulation response to reward applied to spontaneous recording.** Reward distribution for the discrete MAB versus spontaneous activity. The mean is shown as a dashed line. Recorded on DIV 69.

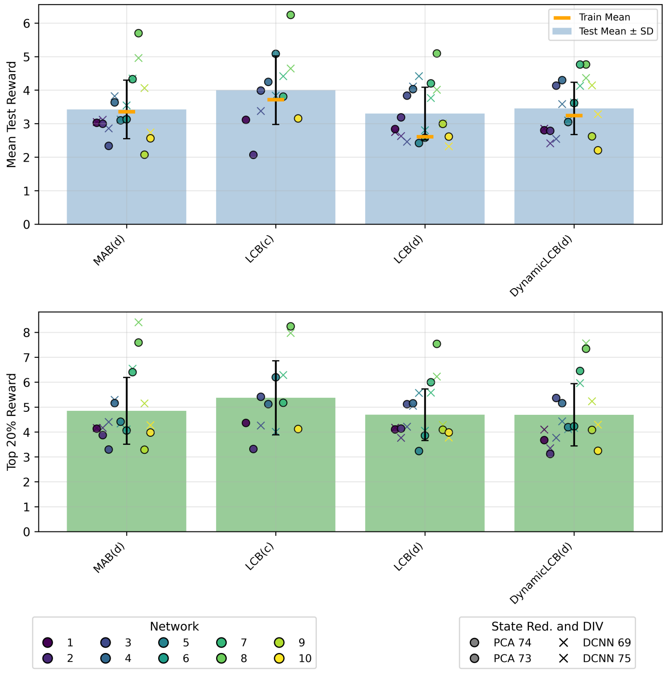


**Fig. S19. Agent performance comparison for PCA and DCNN state compression.** The experiments shown in Fig. 4a,c were repeated with different state compression for one MEA and a subset of agents.

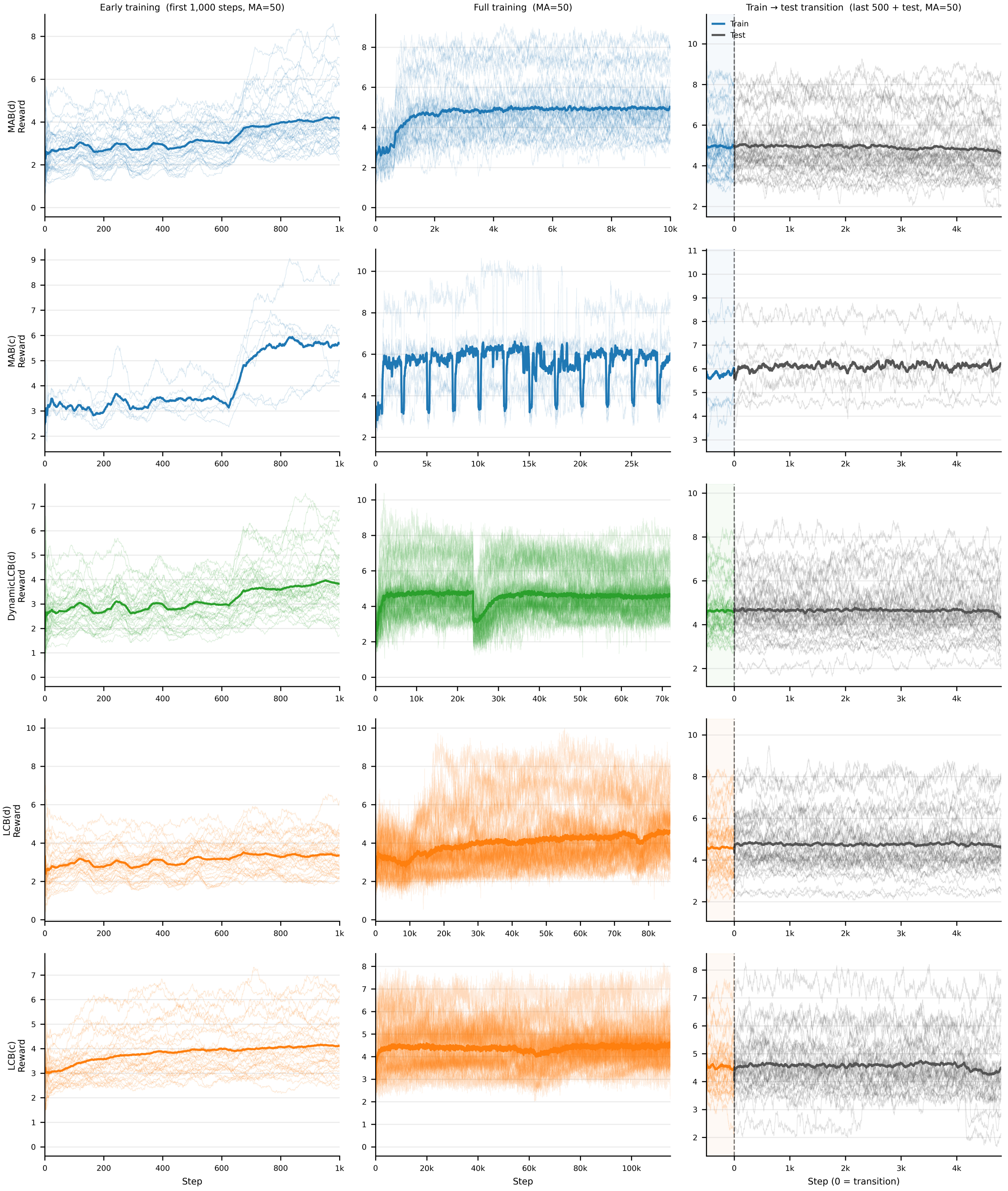


**Fig. S20. Training details for all four agents.** The training data of each agent is shown in a different row. All rewards are filtered with a moving average filter of 50 steps. The first column shows the first 1,000 training steps. The second column shows the full training. The last column shows the transition to the test phase.

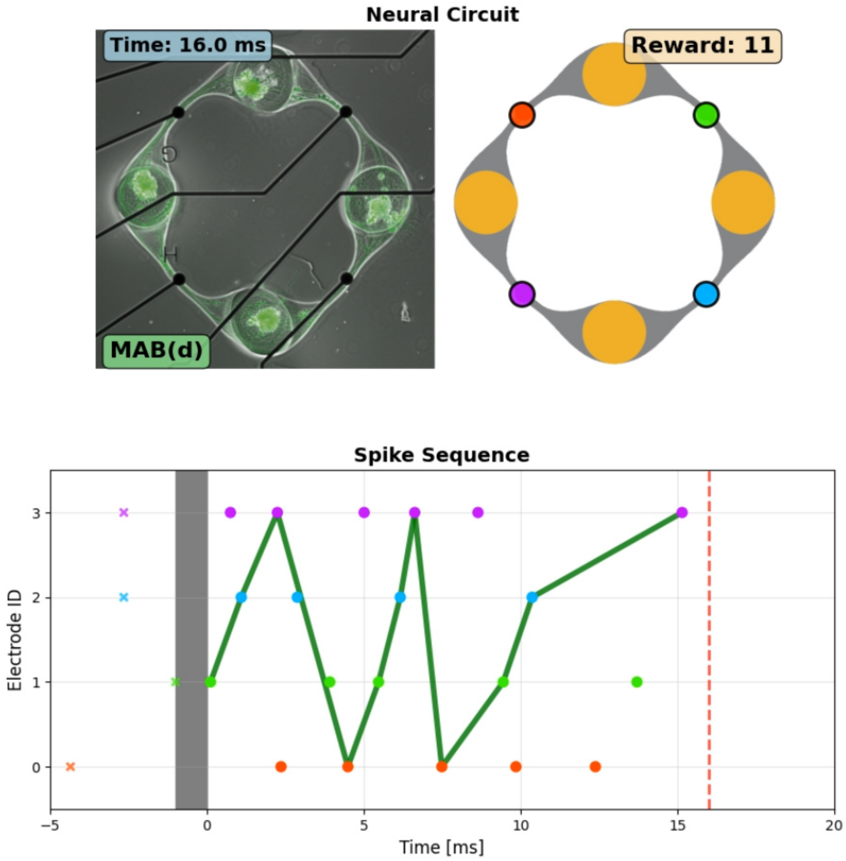


**Fig. S21. Screenshot of a video showing the best-performing test action for each agent.** The video shows the highest rated sequence during the test.

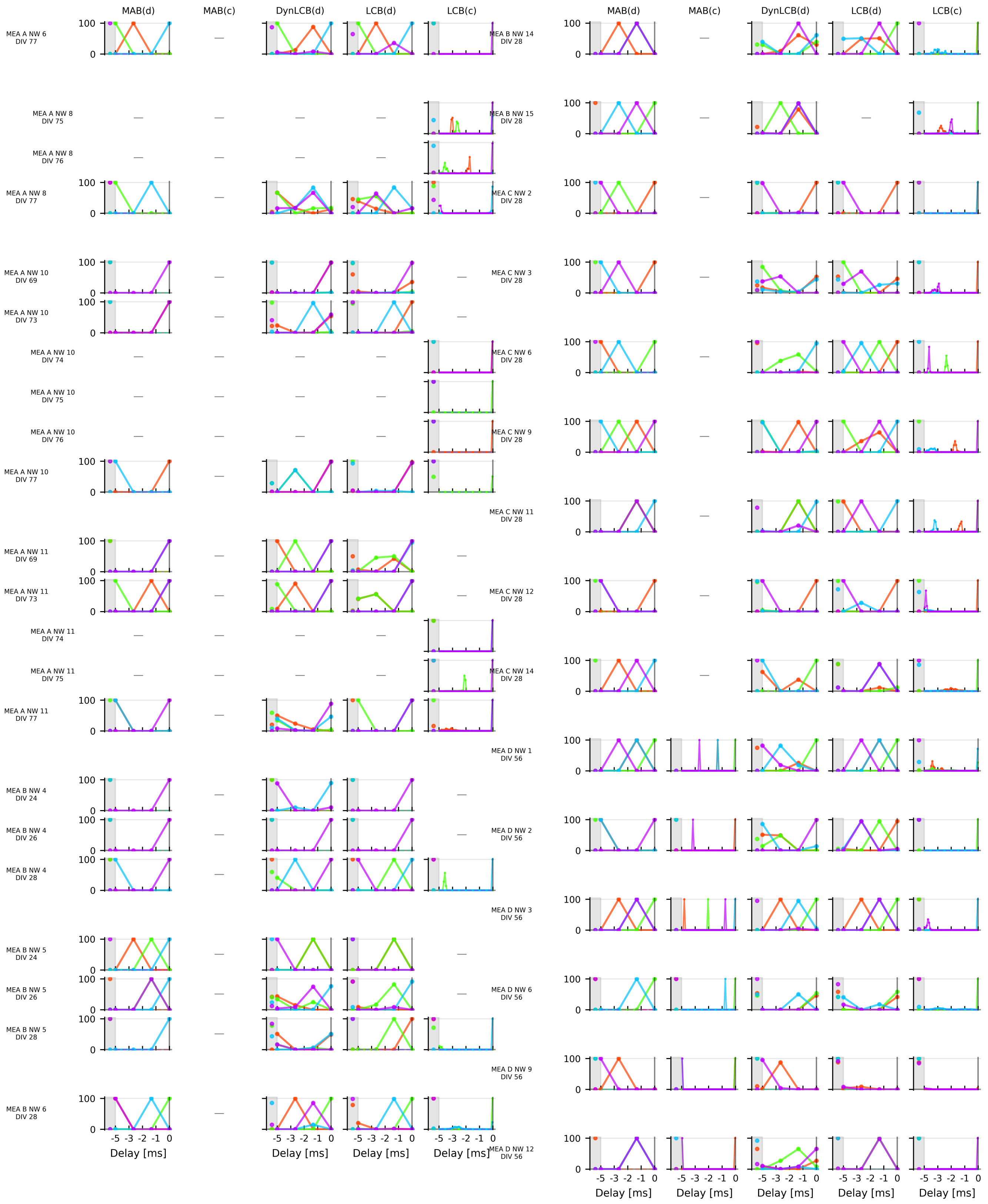


**Fig. S22. Distribution of the test actions across the action space Fig. 5b separated by DIV and network.** The y-axis shows the proportion of test actions in % with the respective action value.
